# Origin and stepwise improvement of vertebrate lungs

**DOI:** 10.1101/2024.07.14.603411

**Authors:** Ye Li, Mingliang Hu, Zhigang Zhang, Baosheng Wu, Jiangmin Zheng, Fenghua Zhang, Jiaqi Hao, Tingfeng Xue, Zhaohong Li, Chenglong Zhu, Lei Zhao, Wenjie Xu, Peidong Xin, Chenguang Feng, Wen Wang, Yilin Zhao, Qiang Qiu, Kun Wang

## Abstract

Lungs, essential for terrestrial vertebrates and present in bony fishes but absent in cartilaginous fishes, provide an ideal model for studying organ origination. Our study analyzed single-cell RNA sequencing data from mature and developing vertebrate lungs, revealing substantial similarities in cell composition, developmental trajectories and gene expression pattern across species. Notably, most lung-related genes are also present in cartilaginous fishes, indicating that gene presence alone does not guarantee lung development. We identified thousands of lung regulatory elements specific to bony fishes, with higher concentrations around genes such as tbx4 and the hoxb gene cluster. These regulatory changes might contribute to lung emergence as well as the unique co-expression patterns in lung epithelial cells, such as those related to pulmonary surfactants and cell morphology. Our research also revealed that AT1 cells are specific to mammals, and we identified a mammal-specific gene, sfta2. Knockout experiments demonstrated that sfta2 deletion causes severe respiratory defects in mice, underscoring its critical role in specialized mammalian lungs. In conclusion, our results demonstrate that the origin and evolution of lungs are driven by a complex interplay of regulatory network modifications and the emergence of new genes, underscoring the multifaceted nature of organ evolution.

## Introduction

While the genesis of new organ systems has been instrumental in major adaptive radiations and ecological transitions throughout life’s history^1–4^, the concrete genetic mechanisms underpinning these remarkable evolutionary innovations remain elusive. In this field, researchers have identified several key processes^4^. One such process involves new genes— whether arising from gene duplication or formed *de novo*—acquiring functions that contribute to the emergence of novel phenotypes. This works in concert with another process where existing genes are repurposed or co-opted, taking on new roles that lead to dramatic phenotypic changes^4–10^. Despite these insights, our understanding of how these mechanisms synergistically operate in the origin of new organs remains limited. This knowledge gap is further exacerbated by the nature of many organs, which are composed of soft tissues that rarely fossilize^11, 12^. This scarcity of fossil evidence makes it challenging to infer intermediate stages in organ evolution and to elucidate the process of their formation.

The evolutionary origin of the vertebrate lung exemplifies these difficulties. Lung is a key innovation enabling vertebrate water-to-land transition. While previous studies have confirmed the homology between fish swim bladders and tetrapod lungs by morphological and genetic evidences^13–16^, the timing and detailed genetic innovation of lung origin remains unclear. Some evidence proposes that lungs may have originated since cartilaginous fishes or ever earlier, as suggested by the presence of a “lung-like” organ observed in Bothriolepis fossils, which represents an outgroup of jawed vertebrate (Gnathostomata), including both bony and cartilaginous species^17^. Moreover, vestiges of potential lungs or air sacs noted in skate embryos at morphological level^18, 19^, as well as lung-specific genetic enhancers near *tbx4* conserved in brown-banded bamboo sharks’ genome^20^, indicate that these enhancers may predate the separation of cartilaginous and bony fish lineages. Conversely, extant adult cartilaginous fish lacking air sacs and the difficulty of preserving chondrichthyan fossils’ soft tissues complicate such interpretations. Additionally, a recent study contesting the identification of the “lung-like” structure in Bothriolepis as a liver suggests that lungs may have evolved as late as the Osteichthyes lineage^12^. Nevertheless, the presence of lungs in bony fish and their absence in cartilaginous fish provide an ideal model for exploring the genetic mechanisms underlying organ origins.

Furthermore, vertebrate lungs exhibit remarkable diversity and specialization in structure and function^21–23^. For example, the simple inflated air sacs with blood vessels in basal ray-finned fish like bichirs contrast with lungfish’s inflated air sacs with folds that enhance their surface area^24, 25^. Furthermore, mammalian lungs’ intricate “respiratory tree” with hundreds of millions of alveoli provides a vast respiratory surface area^26^. Bird lungs, in contrast, have a highly efficient flow-through system in which air flows in one direction, aided by air sacs, allowing continuous and efficient gas exchange during both inhalation and exhalation^21^. Although the morphological structure of many lungs has been extensively described, the molecular mechanisms behind the evolution of complex structures and functions of mammalian lungs from their ancestral counterparts remain elusive.

To investigate the evolution of lung structures, we generated a substantial amount of data, including single-cell transcriptome sequencing for mature and developing lungs across various vertebrates, a new genome assembly of the central bearded dragon, and CUT&TAG sequencing data of chicken lungs. Utilizing a multi-omics approach, we discovered that most lung-related genes were already present in cartilaginous fishes. However, it is the regulatory changes since the common ancestor of bony fishes, rather than new genes, that are key to the presence of lungs. Furthermore, we explored how lung epithelial cells have continued to evolve since their origin, leading to the diverse lung structures observed in extant vertebrates.

## Results and discussions

### The cellular and expression characteristics shared by vertebrate lungs

To trace the origins of lungs, it is crucial to first understand the characteristics of lungs in extant vertebrates. While the lung features of humans and mice are well-documented, these characteristics often include mammal-specific traits that do not represent the entire vertebrate lineage. Thus, to identify shared features across vertebrate lungs, we generated or collected single-cell RNA sequencing (scRNA-seq) data from the lungs of nine species (**Fig. 1a, Supplementary Table 1**). These species include the Senegal bichir (*Polypterus senegalus*), representing the ray-finned fish; the African lungfish (*Protopterus annectens*), representing lobe-finned fish excluding tetrapods; the African bullfrog (*Pyxicephalus adspersus*), representing amphibians; the central bearded dragon (*Pogona vitticeps*), representing reptiles; the chicken (*Gallus gallus*), representing birds; and the pig (*Sus domesticus*), human (*Homo sapiens*), mouse (*Mus musculus*), and rat (*Rattus norvegicus*), representing mammals. The data of first four species were generated in this study (**Extended Data Fig. 1**), while the data for the last five species were collected from previous studies^27, 28^. Collectively, these species effectively cover the major branches of pulmonated vertebrates.

**Figure 1.**
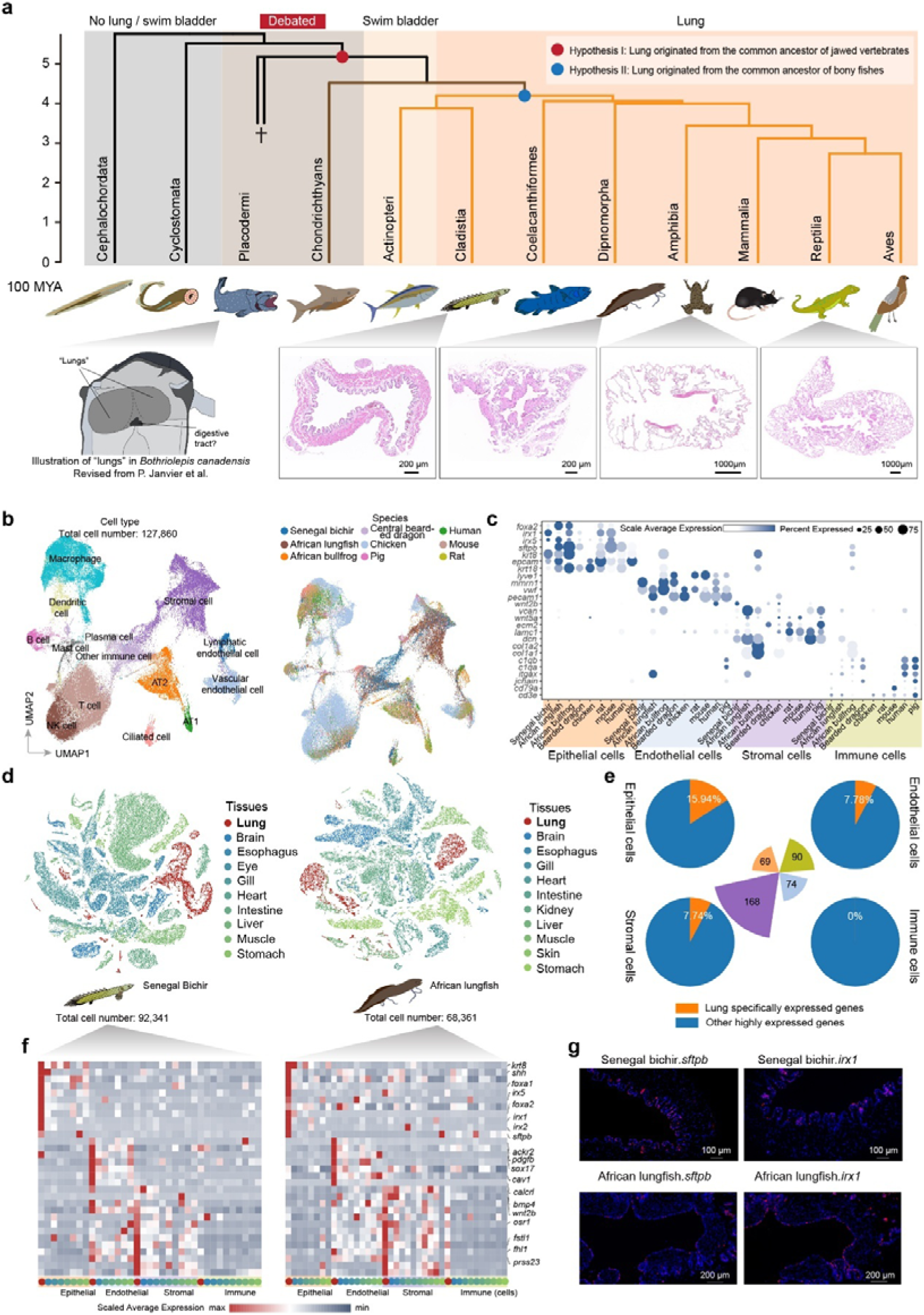
Comparative analysis of vertebrate mature lungs and shared highly expressed genes. **a**, Phylogenetic tree of vertebrates illustrating two hypotheses on lung origin. Red dot: lung originated from common ancestor of jawed vertebrates; Blue dot: lung originated from common ancestor of bony fishes. Bottom panels show schematic and histological sections of lungs across species. **b**, Uniform manifold approximation and projection (UMAP) of integrated lung cells from nine species. Color labels for different cell types (left) and species (right). EP, epithelial; NK, natural killer; AT1, Pulmonary alveolar type I cell; AT2, Pulmonary alveolar type l cell. **c**, Dot plot showing the expression of part of highly expressed genes across four cell types in nine species. Color intensity indicates expression level; dot size represents percentage of cells expressing the gene. **d**, t-distributed stochastic neighbor embedding (t-SNE) plots of cells collected from 10 tissues in adult Senegal bichir (left) and from 11 tissues in adult African lungfish (right). Colors represent different tissues. **e**, Pie charts showing the proportion of lung-specific genes among highly expressed genes in four major cell populations (epithelial, endothelial, stromal, immune cells). **f**, Heatmaps displaying expression of lung-specific genes across cell populations in Senegal bichir (left) and African lungfish (right). **g**, *In situ* hybridization results for *sftpb* and *irx1* genes in lungs of Senegal bichir and African lungfish.

The Canonical Correlation Analysis (CCA) method was applied to integrate 127,860 lung cells from nine species, and the cell populations was annotated using conserved marker genes from humans and mice (**Fig. 1b-c**, and **Extended Data Fig. 2a-b**). To evaluate the robustness of integrating result, we explored alternative methods including SAMAP, scANVI, and four other tools, which, despite variations in integration strength, consistently suggested similar lung cell populations across these species (**Extended Data Fig. 2c**). Combined with the cell populations annotation of each species individually, it can be concluded that epithelium cells (ciliated cells and respiratory epithelial cells), endothelial cells (lymphatic endothelial cells, blood vessel endothelial cells), stromal cells, and immune cells (including lymphoid immune cells such as B cells and T/NK cells, as well as myeloid immune cells like macrophages and monocytes) are shared within these species. Furthermore, we assessed cell-cell communication pathways in lung of each species and discovered that only the VEGF signaling pathway related to vascularity was conserved in all the investigated lungs (**Extended Data Fig. 3a**). This observation suggests that primitive lungs had contained a network of blood vessels for air exchange. It should be noted that while our method specifically highlighted this pathway, the absence of others in our findings does not necessarily imply their non-existence in these species. Additionally, we identified transcription factor (TF) co-expression networks for nine species’ lungs and found 23 conserved TF regulons with high regulatory activity across nine species’ lungs. Among these conserved regulons, the ETS (Erythroblast Transformation Specific) transcription factor family (elf1, elf2, ets1, spi1)^33^, egr1^34^, and klf4^29^ are involved in angiogenesis, while irf1^36^, irf2^37^, and stat1^38^ are associated with immune responses (**Extended Data Fig. 3b**). This finding aligns with the expectation that lungs should be capable of filtering airborne impurities and maintaining immune functions due to their direct contact with air. Collectively, these results underscore the role of primitive lungs as a multifunctional respiratory interface between blood and air, adept at facilitating gas exchange and managing external immune challenges.

To elucidate shared expression characteristics in the lungs at the gene level, we identified 69, 90, 74, and 168 highly expressed genes, respectively (**Supplementary Table 2**), within four major cell populations—epithelial, endothelial, stromal, and immune cells (**Fig. 1c**)— that are highly expressed in more than six out of nine species. Among these, genes highly expressed in epithelial, endothelial, and stromal cells are functionally enriched in tissue morphogenesis, morphogenesis of a branching epithelium, blood vessel morphogenesis, and angiogenesis. This is consistent with the hypothesized structure and function of the primitive lung (**Extended Data Fig. 3c-f**). To account for the inclusion of housekeeping genes typically expressed across various cell types, we also performed single-cell RNA sequencing (scRNA-Seq) on multiple adult tissues from the Senegal bichir and the African lungfish, representatives of ray-finned and lobe-finned fishes, respectively. This additional analysis allowed us to refine our findings and ensure that the identified genes were specifically associated with lung function. In the case of bichir, 92,341 cells and 46 cell types were obtained from 10 tissues including brain, esophagus, eyes, gills, heart, intestine, muscle, stomach, lung, and liver, after filtering out low-quality cells (**Fig. 1d, Supplementary Table 2**). For lungfish, 68,361 cells and 50 cell types were obtained from 11 tissues: brain, esophagus, gills, heart, intestine, muscle, skin, stomach, kidney, lung, and liver (**Fig. 1d**) using the same procedure. From these data, we identified 31 genes (**Fig. 1e**) that showed lung-specific expression in both species, defined as lung-specific genes (using a one-tailed Z-test to determine if their expression in specific lung cell types significantly exceeds that in other cells, p-value < 0.05). Notably, epithelial cells, which serve as the primary parenchymal cells in the lungs, harbored a significant proportion of these lung-specific genes (11 genes, accounting for 15.94% of all highly expressed genes conserved across the nine species). This underscores the specialized role of epithelial cells in the lung, with key genes including *sftpb*, which is essential for lung surfactant function (**Fig. 1f**). The functionally enrichment analysis reveal that these genes linking these genes to structural development and lung morphogenesis, including the development of anatomical structures, tissue morphogenesis, and the morphogenesis of branching and tubular epithelia (**Fig. 1g**), which are associated with lung’s self-renewal and maintenance processes^30^. Additionally, among the 13 lung-specific genes identified in stromal cells, there are key developmental genes such as *osr1*, *wnt2b* and *bmp4*. Although traditionally associated with developmental roles, their expression in the adult lung suggests a potential involvement in maintaining lung tissue architecture and responding to pulmonary stress or injury.

We further examined the gene expression level of these 31 lung-specific genes in the single cell atlas of mouse (**Extended Data Fig. 3g**) and found some of them, including *Sftpb*, *Irx1*, *Irx5*, *calcrl* and *fhl1*, are showing lung-specific expression pattern, but many other genes are also expressed in multiple tissues. Therefore, due to differences in the organs among various species, tissue-specific expression genes may vary significantly between species^31^, in our subsequent exploration of the origin of lungs, we primarily utilized the set of genes that are highly expressed in the lungs of most species.

### Conserved lung developmental genes and signaling pathways

The above analysis has identified a set of highly expressed genes shared by vertebrate’s mature lungs, while a distinct set of genes are employed during lung development. To explore the genes and cellular developmental pathways shared in lung development across different species, we gathered published single-cell transcriptomic data from various stages of mouse lung development (E12, E15, E18, P0, P3, P5, P7 and P14), comprising a total of 100,546 cells^32^ (**Extended Data Fig. 4a**). Furthermore, we performed single-cell sampling from chickens at three developmental stages: Embryonic Day 6 (E6) and Embryonic Day7 (E7), which are respectively the sixth and ninth days after fertilization, and Postnatal Day 3 (P3), occurring three days post-hatching. After filtering out low-quality cells and blood cells, we obtained a total of 39,152 cells. Additionally, we collected published single-cell data from chickens on P32, representing adult stages, amassing a total of 86,155 cells for further analysis (**Fig**. **2a-c**). Given that chicken and mouse diverged approximately 319 million years ago, the shared expression traits between these two species could partially represent the core vertebrate lung developmental pathways.

**Figure 2.**
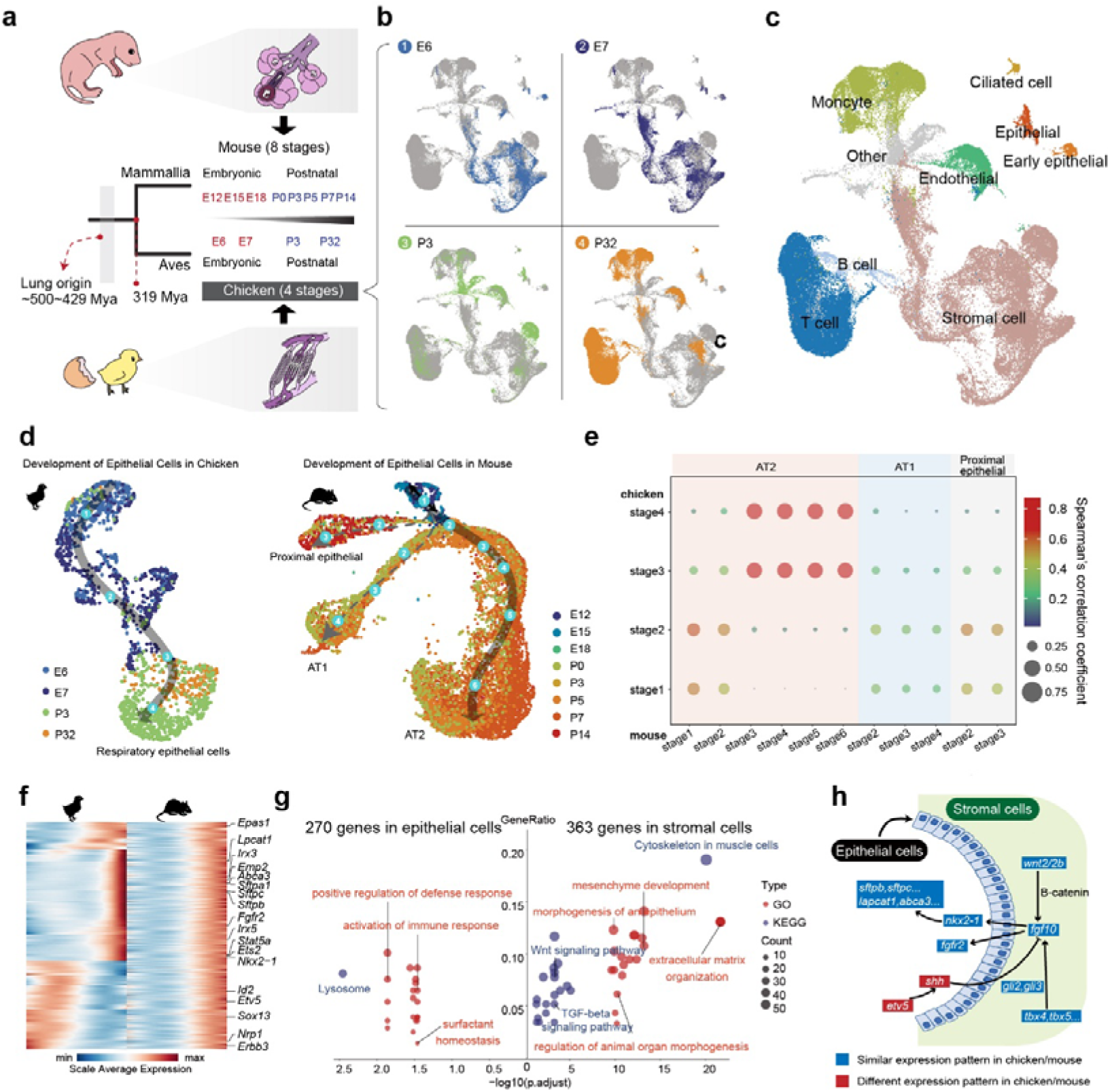
Comparative analysis of developing lungs in mouse and chicken. **a**, Evolutionary timeline and experimental design for scRNA-seq analysis of lung development in mouse (8 stages) and chicken (4 stages), including embryonic and postnatal time points. **b**, UMAP plots of chicken lung cells at four developmental stages (E6, E7, P3, P32). **c**, UMAP plot showing major cell types identified in developing chicken lung. **d**, Developmental trajectories of epithelial cells in chicken (left) and mouse (right). Colors represent different developmental stages. **e**, Dot plot comparing correlation of epithelial cell stages between chicken and mouse. Color intensity indicates Spearman’s correlation coefficient; dot size represents significance. **f**, Heatmaps displaying gene expression patterns across pseudotime in chicken respiratory epithelial cells (left) and mouse AT2 cells (right). **g**, Bubble plot showing enriched GO terms and KEGG pathways in epithelial (270 genes) and stromal (363 genes) cells. Bubble size indicates gene count; color represents cell type. **h**, Schematic illustrating key gene interactions between stromal and epithelial cells during lung development. Colors indicate similar (blue) or different (red) expression patterns between chicken and mouse.

Given that respiratory epithelial cells are highly specialized for the primary function of the lungs - facilitating gas exchange - we first examine the developmental trajectory of epithelial cells in the lungs of chickens and mice. Utilizing trajectory inference tools^33^, we mapped the progression of mouse lung epithelial progenitor cells into three distinct lineages: proximal epithelial cells (primarily secretory cells), and distal alveolar type 1 (AT1) and alveolar type 2 (AT2) cells (**Fig. 2d**). In chickens, we identified a single trajectory from progenitor to respiratory epithelial cells (**Fig. 2d**). Importantly, the expression patterns of non-mammalian respiratory epithelial cells exhibit high similarity to mammalian AT2 cells, and their progenitor cells in both species also share similar expression profiles (**Fig. 2e**). This suggests that the developmental pathway from lung epithelial progenitor cells to respiratory epithelial cells in chickens, or to AT2 cells in mice, represents a homologous developmental route and a key pathway shared by vertebrate lung epithelial cells. Our analysis revealed 270 genes exhibiting a progressive increase in expression levels in both species, with significant overlap (P-value = 4.78e-51), underscoring the homology of their developmental trajectories (**Fig. 2f, Supplementary Table 2**). These genes are enriched in functions related to lipid synthesis and metabolism, surfactant metabolism, and immune responses (**Fig. 2g**). Notably, many lung morphogenesis genes are sharing similar expression pattern in the developmental pathways of chicken and mouse (**Fig. 2h, Extended Data Fig. 4b**), including *nkx2-1*, the master regulator essential for lung development, differentiation, and maintenance^34^. Similarly, the downstream genes for lung development activated by *nkx2-1*, such as the sftp gene family, the genes (*abca3*, *slc34a2*) involved in the transport and secretion of pulmonary surfactant, the synthesis and supply of surfactant materials, are also shared by two species. Additionally, we observed genes gradually upregulated in mouse lung epithelium, but not in chickens. Some are mammal-specific highly expressed genes (**Extended Data Fig. 4c**).

Furthermore, previous studies have emphasized the crucial role of stromal cell signals in lung morphogenesis and epithelial cell differentiation. To identify conserved stromal signals critical for lung development across species, we analyzed the interaction pathways between stromal and respiratory epithelial cells in both chicken and mouse. Our analysis revealed 32 shared signaling pathways in stromal cells (**Extended Data Fig. 4d-e**), including WNTs, BMPs, and TGFβ. Consistently, we identified 363 genes highly expressed in stromal cells of both mouse and chicken developing lungs, which are enriched in several important signaling pathways (**Supplementary Table 2**). Among these, the *fgf10* gene is known to activate *nkx2-1* in epithelial cells^35^ (**Fig. 2h**). Additionally, genes that promote the expression of *fgf10*, such as *tbx4/5* and *wnt2/2b*, are also highly expressed in the stromal cells of chicken and mouse embryos (**Extended Data Fig. 4f**). Our analysis further uncovered intriguing similarities and differences in gene expression patterns between mouse and chicken epithelial cells interacting with stromal cells. While some genes, such as *fgfr2* (a receptor for Fgf10 that plays a crucial role in regulating precise lung branching in epithelial cells^36^), show conserved expression patterns across both species, we identified two important genes with divergent expression patterns (**Fig. 2h**). The first is *shh* (Sonic hedgehog), which inhibits Fgf10 and controls lung branching morphogenesis^37^; it is expressed at relatively higher levels in the later stages (AT1) in mice, while in chickens its expression peaks in the middle stages of development (**Extended Data Fig. 4b**). The second gene is *etv5*, which primarily regulates the periodicity of lung branching morphogenesis^38^; it shows high expression in the later stages of mouse lung development but is highly expressed in the early stages in chickens (**Fig. 2f, Extended Data Fig. 4b**). These differences in expression patterns suggest potential species-specific mechanisms in lung development between mice and chickens.

### Evolutionary dynamics of genes and enhancers required by mature and developing lungs

Collectively, the above analysis has identified a set of 881 lung-related genes. These genes are either highly expressed in most vertebrate lungs, specifically 327 genes in epithelial cells, endothelial cells, and stromal cells (excluding those highly expressed in immune cells because their gene expression patterns are highly complex and variable, depending on different stages of immune responses), or play significant roles during lung development, accounting for 631 genes. The next step is to determine when these genes originated and whether their origins provide insights into the evolution of lungs. Remarkably, the vast majority of these genes (860) originated in cartilaginous fish or even earlier in evolutionary history (**Fig. 3a, Supplementary Table 2**). In contrast, only six genes (*areg*, *ctsk*, *dcun1d2*, *matn2*, *ac142391.1* and *cd47*) were emerged in the lineage leading to the common ancestor of bony fishes. Notably, more than one-third of these lung-related genes are products of the two rounds of whole genome duplication (2R-WGDs) shared by jawed vertebrates, a proportion significantly higher than the whole genome background (**Fig. 3b**). This group includes genes essential for lung function and development, such as *sftpb* (**Extended Data Fig. 5a**), *foxa2*, *wnt2b*, and *shh*. This finding highlights the ancient origins of most lung-related genes and underscores the significant impact of 2R-WGDs on the evolution of complex organs^39^.

**Figure 3.**
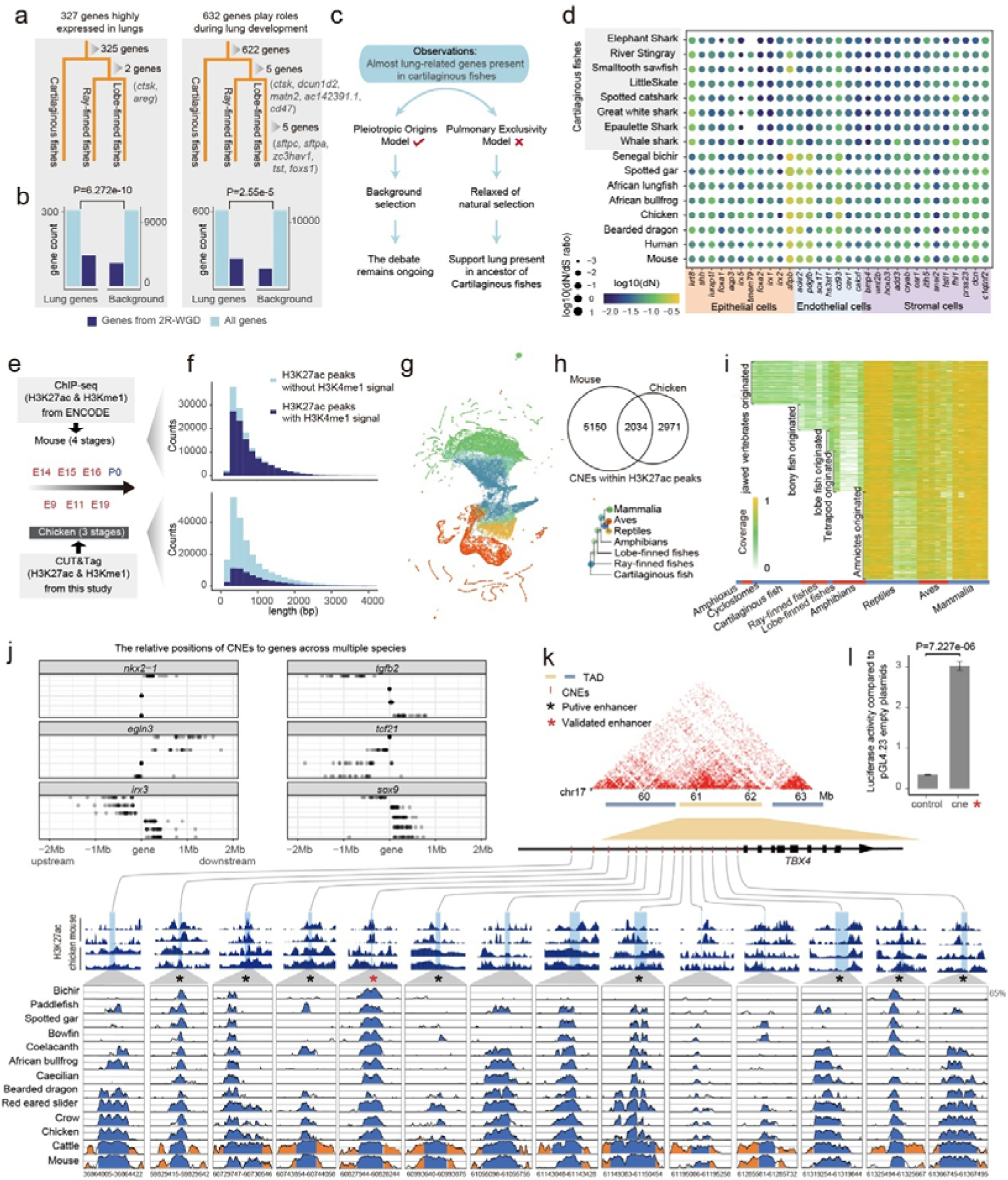
The evolution of lung-related genes and regulatory elements in cartilaginous and bony fishes. **a,** Phylogenetic origin of 327 genes highly expressed in mature lungs and 632 genes involved in lung development. Genes with more recent origins after jawed vertebrates emerged are listed. The tree shows the evolutionary relationships and gene counts at each branch. **b,** Comparison of 2R-WGD (two-rounds of whole genome duplication) gene ratios between lung-related genes and background genes. Bar graphs show gene counts for lung genes and background genes from 2R-WGD and all genes. **c,** Observations on lung gene origins and two explanatory models: the pleiotropic origins model (supported) and pulmonary exclusivity model (rejected). The latter is dismissed due to lack of relaxed natural selection signals for lung-related genes in cartilaginous fishes. **d,** Heatmap showing the presence of partial lung-related genes across various species, from elephant shark to human. Color intensity indicates log10(N) of gene counts. **e,** Schematic of ChIP-seq and CUT&Tag data used for mouse (4 stages) and chicken (3 stages), including newly generated chicken CUT&Tag data in this study. **f,** Length distribution of H3K27ac histone binding peaks in mouse and chicken. Dark blue represents peaks with both H3K27ac and H3K4me1, light blue represents H3K27ac-only peaks. **g,** UMAP visualization of regulatory elements active in embryonic mouse and chicken lungs, based on alignment percentage to 45 vertebrate genomes. Colors represent different taxonomic groups. **h,** Venn diagram showing overlap of amniote-conserved elements with lung activity in mouse and chicken. Numbers indicate shared and unique elements. **i,** Evolutionary origin of amniote-conserved elements across species groups. Color intensity indicates alignment coverage, with yellow representing higher coverage. **j,** Relative positions of some lung-related genes (*nkx2-1*, *wnt2*, *agr2*, *igf21*, *fgf3*, and *spp1*) and nearby CNEs (Conserved Non-coding Elements) across species. Each row represents a CNE, with dots indicating its position relative to the gene in different species. **k,** Detailed view of 14 CNEs near the *TBX4* gene. Top: human lung Hi-C interactions, TAD (Topologically Associating Domain) distribution, and CNE positions (using hg38 as reference). Middle: H3K27ac signals in chicken and mouse embryonic lungs. Bottom: Sequence conservation across multiple species. Asterisks indicate CNEs with enhancer signals, with red asterisks denoting experimentally validated CNEs. **l,** Bar graph showing dual-luciferase assay results, demonstrating >9-fold enhancer activity for the red-asterisked CNE compared to control.

Reconsidering the debates about lung origin, we question whether lungs originated from cartilaginous fish or bony fish (**Fig. 3c**). Intuitively, the presence of these genes in cartilaginous fish seems to support the hypothesis that lungs originated from them. However, it is possible that the emergence of these genes did not necessarily lead to the development of lungs but rather initially served other functions during vertebrate evolution (Pleiotropic Origins Model). An extreme counter-model posits that these genes originally appeared specifically for lung development (Pulmonary Exclusivity Model). If this were true, we would expect to see relaxed selection in some cartilaginous fish, given that lungs are absent in them. To test this hypothesis, we collected the orthologs of all lung-related genes in both eight cartilaginous and eight bony fish species, calculated their nonsynonymous/synonymous (dN/dS) ratio, and observed no significant difference between the two groups (**Extended Data Fig. 5b**). This seems to disprove the extreme counter-hypothesis, indicating that even if cartilaginous fish had lungs, these genes were not exclusively for lung development. While we cannot definitively determine whether lungs originated in cartilaginous fish, these results demonstrate that the presence or absence of lung-related genes is not the decisive factor in determining lung existence. This observation aligns with a well-established hypothesis in evolutionary biology: that changes in gene regulation, rather than the emergence of new genes, often drive the development of novel traits. Consequently, it is likely that regulatory elements unique to bony fish played a crucial role in the origin and evolution of lungs.

Our investigation then focused on the regulatory elements crucial to lung function and development. Due to the spatiotemporal specificity of regulatory elements and their crucial role in organ development, we collected enhancer sequences from mouse and chicken lung development to explore their evolution in vertebrates (**Fig. 3e**). First, we conducted CUT&Tag with histone H3K27ac and H3K4me1 antibody for day 9, 11, and 19 chicken lungs, and identified 62.66 Mb (with a median length of 573 bp) regions with H3K27ac activity in chicken lung development, indicating their regulatory roles, and 52.63% of these regions are also with H3K4me1 activity, indicating their role of enhancer (**Fig. 3f, Extended Data Fig. 5c**). Meanwhile, we identified 84.74 Mb (with a median length of 608 bp) regions with H3K27ac activity during mouse lung development using related ChIP-seq data from ENCODE project^40^, and 54.50% of these regions are also with H3K4me1 activity (**Fig. 3f**). These sequences were then mapped to 48 vertebrate species to illustrate their evolutionary trajectory. The result showing that while there is a large proportion of regions with enhancer activity are lineage specific to mammalians or avians, there are also about 51.7% of all sequences shared by amniotes and even have an earlier origin history, reflecting the shared regulatory basis for lung development across species (**Fig. 3g**).

Since the sequences with enhancer activity in mouse or chicken are quite long and are not conserved across species as a whole, we extracted the conserved non-coding elements (CNEs) within these regions across 21 amniotes species. These CNEs were then used as the basic units to study the origin of lung-development-related regulatory elements. In total, we identified 10,155 CNEs with regulatory activity in either mouse or chicken (**Fig. 3h**). Within them, 40.64% of the CNEs that exhibit regulatory activity in chickens also show enhancer activity in mice. Using GREAT, we performed the functional enrichment of adjacent genes close these CNEs, and found they are mainly associated with cell proliferation and differentiation, including lung cell differentiation and lung morphogenesis (**Extended Data Fig. 5d**). By further determining the origin nodes of these CNEs, we identified a total of 1,162 CNEs originating from the common ancestor of bony fishes and 2,836 CNEs originating from the common ancestor of jawed vertebrates or even earlier (**Fig. 3i, Supplementary Table 3**). To determine which bony fish CNEs that exhibit regulatory activity during lung development are associated with the recruitment of lung highly expressed genes or developmental genes, we mapped these CNEs across 35 representative bony fish species. We searched for gene-CNE pairs that are in close proximity (within in 2 Mb) in more than 18 species, as the conservation of relative positions suggests a functional association. In total, we identified 913 such gene-CNE pairs (**Supplementary Table 4**), comprising 369 lung-related genes and 561 CNEs, including well-established genes critical for lung development, such as *Nkx2-1* (3 associated CNEs).

Notably, these gene-CNE pairs are not evenly distributed around the 375 genes. A subset of 16 genes accounted for 192 gene-CNE pairs, each associated with 10 or more CNEs. *Tbx4*, crucial for lung development, stands out with 14 gene-CNE pairs (**Fig. 3k**). This gene regulates FGF10 expression, a key growth factor for lung bud formation and branching morphogenesis^41^. All 14 CNEs associated with *tbx4* are located upstream of the gene, with 10 showing clear enhancer signals. Human lung Hi-C data reveals that 12 of these 14 CNEs and *tbx4* reside within the same topologically associating domain (TAD), suggesting potential functional interactions. To validate the function of these CNEs, we also performed dual-luciferase assays on one of them using human alveolar basal epithelial cells (A549), and confirmed the enhancer activity of the tested element (**Fig. 3l**). Some CNEs cluster around specific gene groups. For example, the *Hoxb* gene cluster, including *Hoxb3*, *Hoxb5*, *Hoxb6*, and *Hoxb8*, has 10 nearby CNEs, of which 5 exhibit clear enhancer signals (**Extended Data Fig. 6**). Expression data indicate that these *Hoxb* genes are predominantly expressed in stromal cells, suggesting their potential role as signaling molecules influencing lung branching and epithelial cell differentiation. This clustering distribution of CNEs suggests that lung formation requires a complex regulatory program. Moreover, this intricate regulatory network, unique to bony fish, might be a key determinant for the presence of lungs in these species.

To further identify which transcription factors (TFs) regulate these CNEs, we scanned for their binding sites (TFBS), identifying 936 relevant TFs (**Supplementary Table 5**). Some TFs interact with a large number of CNEs, while a few interact with only one CNE. The TF with the most connections is VEZF1, with its 375 binding sites found in 266 CNEs. Previous studies have shown that VEZF1 plays a crucial role in the formation of the vascular system and is broadly expressed in various tissues and cell types^42^. The widespread presence of VEZF1’s biding sites, originating from bony fish and accessible in the chromatin of the lung cells, suggests that VEZF1 may facilitate the development of the pulmonary vascular network through a complex global regulatory mechanism. For instance, several transcription factors highly expressed in lung epithelial cells also interact with numerous CNEs. Notably, FOXA1 and FOXA2 are connected to 50 and 47 CNEs, respectively, including CNEs near *tbx4*. These interactions suggest that genes in the lung work as a complex network to collectively promote lung formation and functional maintenance.

Overall, the conservation of gene-CNE pairs across multiple bony fish species and their uneven distributions suggest a complex regulatory network essential for lung development. It is important to note that our analysis is limited to conserved elements with H3K27ac histone signals, and many regulatory elements may not be sequence-conserved. Therefore, the extent of regulatory changes since the evolution of bony fishes is likely more substantial than our current findings suggest.

### Respiratory epithelial cell evolution from chondrichthyans to terrestrial vertebrates

While our findings demonstrated that most lung-related genes are already present in chondrichthyans, it is interesting to explore how these lung-specific genes are expressed in cartilaginous fishes that without lungs. Therefore, we sequenced scRNA-seq transcriptomes from 14 organs, along with bulk transcriptomes from 19 organs of white spotted bamboo shark (*Chiloscyllium plagiosum*) (**Fig. 4a**). A total of 107, 652 cells were obtained after filtering and 55 cell types were obtained after clustering and annotation (**Fig. 4a**). While lung-specific genes are broadly expressed across different cell types in the bamboo shark, they do not exhibit tissue specificity (**Fig. 4b**). The expression matrix from bulk transcriptome data shows a similar pattern (**Extended Data Fig. 7a**). These results highlight the pleiotropy of lung-specific genes in the bamboo shark. For example, the gene *sftpb* is broadly expressed in many organs (with an average FPKM of 84.9 in bulk RNA-seq) of bamboo shark (**Extended Data Fig. 7b**). Notably, *sftpb* is highly expressed in the spleen and specifically expressed in immune cells in some organs. *In situ* expression experiments confirmed that *sftpb* is indeed expressed in the spleen, liver, and gill, but not in muscle (**Fig. 4b**). Given that both the spleen and liver are crucial for immunization and the gill is directly exposed to the external environment, it is likely that *sftpb* primarily plays an antibacterial or immunoregulatory role^43^ in many organs of bamboo shark. Similar to *sftpb*, other lung-specific genes are also expressed in various tissues of bamboo shark. Overall, the dispersed expression of these genes in various tissues of bamboo shark, along with their specifically high expression in the epithelial cells of the lung, further exemplifies that gene recruitment is a key factor in the emergence of lung epithelial cells as an innovation in bony fish.

**Figure 4.**
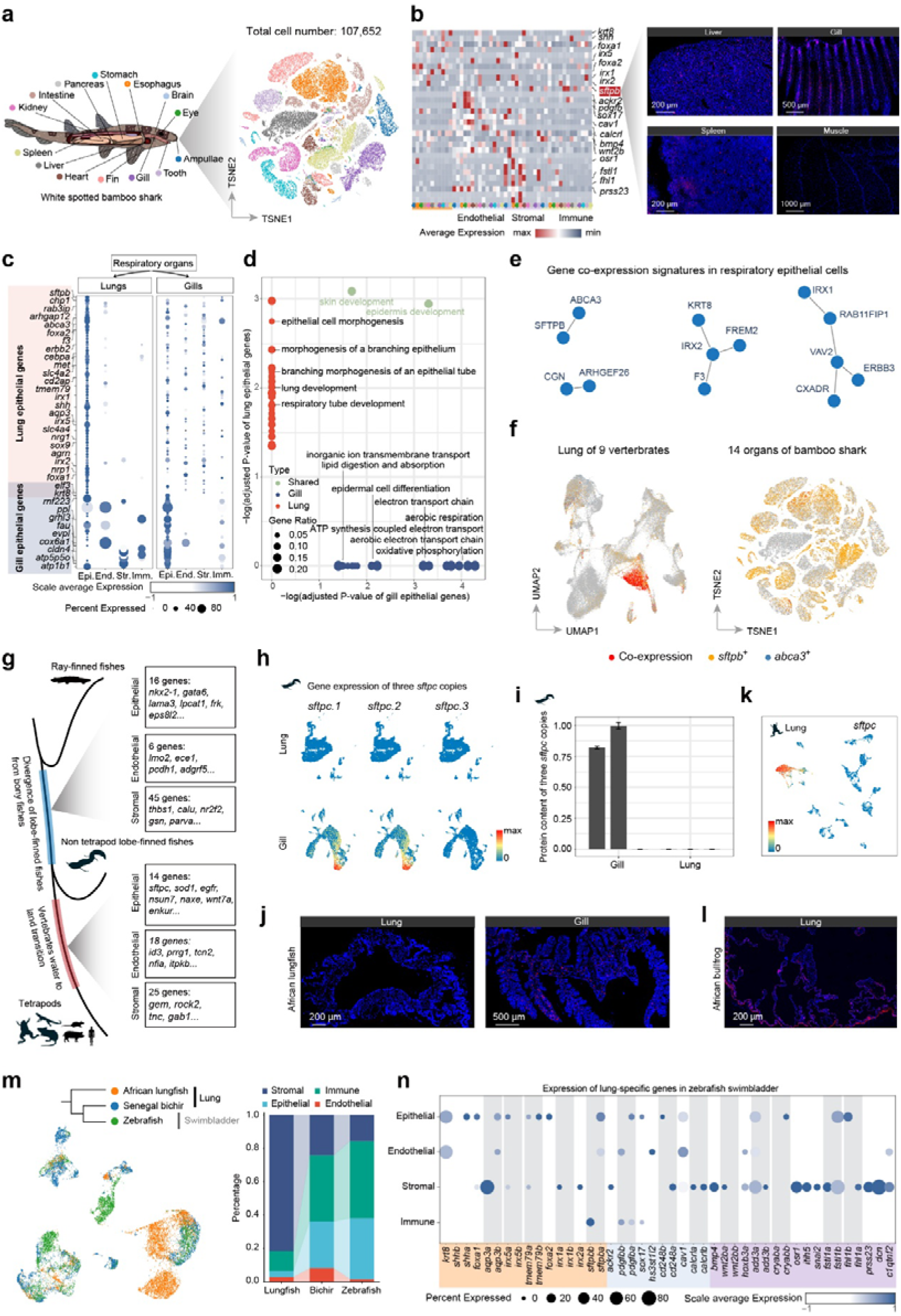
Expression basis and evolutionary changes for respiratory epithelial cell in vertebrate evolution. **a,** t-SNE plot of single-cell transcriptomic profiles from 14 tissues in adult white spotted bamboo shark (107,652 total cells). Cells are colored according to tissue origin. **b,** Left: Heatmap displaying average expression of lung-specific genes across cell populations in white spotted bamboo shark. Right: *In situ* hybridization analysis of *sftpb* in four tissues (liver, gill, spleen, muscle) of the white spotted bamboo shark. **c,** Dot plot showing expression of conserved genes highly expressed in gill or lung epithelial cells. **d,** Bubble plot displaying enriched GO terms with genes conserved and highly expressed in lungs and gills for epithelial cells. **e,** Network diagram of genes with significantly higher co-expression in lung epithelial cells. **f,** UMAP and tSNE plots showing co-expression of *sftpb* and *abca3* in lungs of 9 vertebrates (left) and 14 organs of bamboo shark (right). **g,** Schematic diagram showing genes highly expressed in lobe-finned fish and terrestrial vertebrates across different cell populations (epithelial, endothelial, stromal). **h,** Feature plots displaying expression of three *sftpc* copies in gill and lung of the African lungfish. **i,** Bar graph showing relative content of *sftpc* copies in gill and lung of the African lungfish from proteome sequencing data. **j,** *In situ* hybridization results showing expression of *sftpc* (*s*ftpc.1) in gill and lung of the African lungfish. **k,** Feature plot displaying expression of *sftpc* in the lung of African bullfrog. **l,** *In situ* hybridization results showing expression of *sftpc* in the lung of African bullfrog. **m,** Left: UMAP plot showing integrated cell clustering for lungs of African lungfish, Senegal bichir, and swim bladder of zebrafish. Right: Bar chart showing proportion of each cell population (stromal, immune, epithelial, endothelial) in these three species. **n,** Dot plot displaying expression of lung-specific genes in different cell populations of the zebrafish swim bladder. Gray shaded areas highlight genes with two retained copies post whole-genome duplication.

We then compare the lung epithelial cells with gill epithelial cells, another representative respiratory organ. Our results show that only two (*krt8* and *elf*) of the 69 genes shared by lungs of nine bony fish species overlap with the 25 genes shared by gills of three species (bamboo shark, bichir and lungfish) (**Fig. 4c**). Despite both being respiratory organs, the differentially expressed genes highlight the significant differences required for their distinct respiratory surfaces. In lung epithelial cells, highly expressed genes are enriched in functions associated with epithelial tube morphogenesis and branching (**Fig. 4d**). In contrast, gill epithelial cells exhibit gene enrichment in functions related to aerobic respiration, electron transport chain, and inorganic ion transmembrane transport (**Fig. 4d**). This suggests that the respiratory epithelium in gills requires high expression of genes facilitating ion transport to maintain osmotic pressure in an aquatic environment, which also demands substantial energy supply. Conversely, lung respiratory epithelium primarily needs to maintain a structure conducive to the free diffusion of air molecules.

It is hypothesized that new cell type is evolved with the emergence of new protein and protein interaction^44^. To explore potential new interactions in lung respiratory epithelial cells, we identified 9 gene pairs with significantly higher co-expression in lung respiratory epithelial cells across nine vertebrate species compared to the 55 cell types in bamboo shark (**Fig. 4e, Supplementary Table 6**). These gene pairs showed at least 5-fold higher co-expression ratios in lung epithelial cells (|Z-score| > 1.65) relative to the maximum ratio in shark cell types, suggesting their potential involvement in lung functions. These gene pairs can be functionally categorized into two main groups, reflecting key aspects of lung epithelial cell biology. The first group, comprising *abca3* and *sftpb* (**Fig. 4f**), is directly involved in surfactant-related functions, with *abca3* responsible for surfactant transport and *sftpb* serving as a crucial component of lung surfactant^45^. These two genes exhibit a high degree of co-expression in lung epithelial cells, while their co-expression levels are notably low in any cell type in bamboo shark. This observation was further corroborated by bulk transcriptome analysis (**Extended Data Fig. 7b**). The second group, encompassing the remaining gene pairs, is primarily associated with cellular structure, intercellular connections, and cell polarity. These genes are involved in various processes including cell adhesion, signal transduction, cytoskeleton regulation, membrane protein transport, and epithelial differentiation. This co-expression pattern suggests that lung epithelial cells may have evolved specialized regulatory networks or Core Regulatory Complexes (CoRCs)^44^ to support both lung-specific functions (surfactant-related) and modified epithelial functions (cell structure and organization). Additionally, we also observed several of lung regulatory elements unique to bony fish around some of these genes, such as *vav2* (5 related CNEs), *cxadr* (7 related CNEs), *cgn* (4 related CNEs) and *irx1*/*irx2* (two shared related CNEs), which might have contributed to their co-expression in lung epithelial cells.

Since the expression pattern lung epithelial cells provide an important basis for air respiration, then what contribute to vertebrate water-to-land transition? We compared the single cell transcriptome data between African lungfish, Senegal bichir and tetrapods. While no significant divergence in cell composition, there are an increased number of conserved expressed genes in the lungs of lobe-finned fishes (African lungfish and tetrapods) and in the lungs of tetrapods (**Fig. 4g**). The lung conserved in lobe-finned fishes are including *Nkx2-1* and the lung surfactant gene *Lpcat1*. These genes are enriched in functions associated with alveolar development and angiogenesis (**Extended Data Fig. 7c**). The lung conserved genes in tetrapods are including *Sftpc*, which is originated since the LCA of lungfish and tetrapods and have three copies in lungfish. While all the three copies have low expression level in the lungs of African lungfish, two of them are highly expressed in its gills (**Fig. 4h**). To validate this observation at protein level, we also generated proteomic data for gills and lungs of African lungfish, and the results suggest that two copies of *Sftpc* are indeed detected in gill but not lung (**Fig. 4i**). In contrast, *Sftpc* is highly expressed in all the seven tetrapod lungs, and is barely expressed in frog’s tadpole gills^46^. This pattern suggests that *sftpc* originally did not initially used for lung function, but later was recruited to have a vital role in tetrapod lungs.

As a significant branch of bony fish, teleost fish have undergone an independent genome duplication. Unlike lungs, most teleost swim bladders lack respiratory function. To study the expression of lung-specific genes in the swim bladder, we conducted single-cell sequencing on zebrafish swim bladders, obtaining 4,171 cells after filtering out red blood cells (**Fig. 4m**). After annotating these cells, we examined the expression of lung-specific genes in swim bladder cells (**Fig. 4n**). In epithelial and stromal cells, certain lung-specific genes were expressed consistently in the corresponding cell types of the swim bladder, such as *foxa2* and *thmem79*. Following whole-genome duplication, the expression of some gene copies changed; for example, *s*ftpba is expressed in epithelial cells, while *s*ftpbb is expressed in immune cells. Additionally, some genes retained their original expression pattern in one copy while the other copy was not expressed, such as *irx5*.

### Lineage-specific specialization in mammalian lungs

In contrast to other vertebrates, mammalian lungs exhibit significantly enhanced respiratory efficiency through the reduction of alveolar size. To elucidate the unique cellular composition and gene expression profiles in mammalian lungs, we conducted a comparative analysis of single-cell transcriptome data from mammalian lungs and those of other species, including Senegal bichir, African lungfish, African bullfrog, and central bearded dragon (**Fig. 5a**). Our analysis revealed that mammalian lungs possess a greater diversity of cell types, including further differentiation of endothelial cells (**Extended Data Fig. 8a-c**). Among these specializations, the most significant finding was the distinct categorization of mammalian epithelial cells into AT1 or AT2 types, a differentiation not observed in non-mammalian species (**Fig. 5b**). To validate this observation, we performed *in situ* expression analysis targeting *clic5* (AT1 marker) and *sftpb* (AT2 marker) in the lungs of mice, central bearded dragons, and African bullfrogs (Fig. 5c). Results demonstrated high and adjacent expression of both *clic5* and *sftpb* in mouse lungs. Conversely, we observed predominant *sftpb* expression in the lungs of central bearded dragons and African bullfrogs, with *clic5* expression being notably diminished, particularly in African bullfrogs where it was nearly absent.

**Figure 5.**
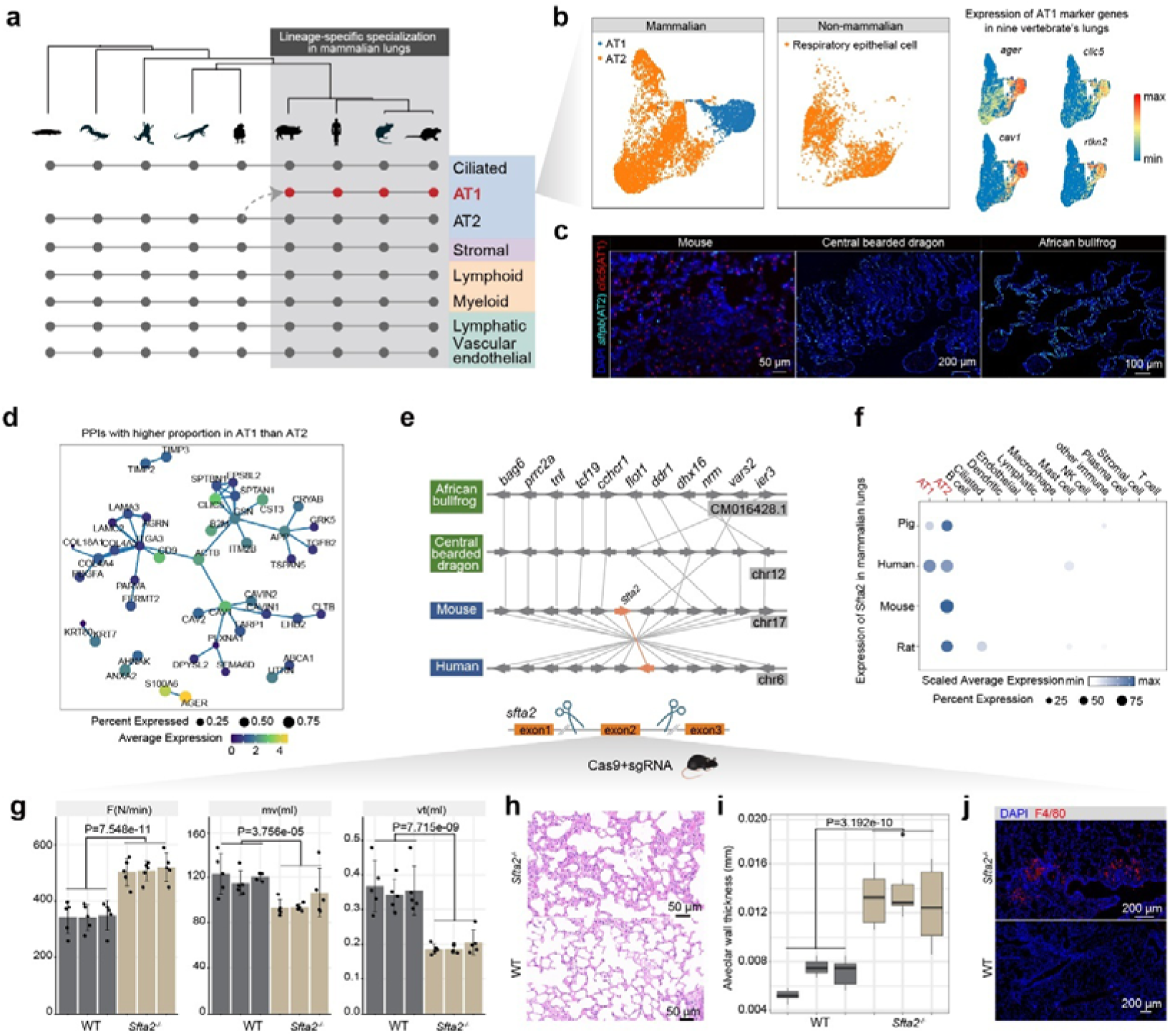
The emergence of a new cell type (AT1) and gene (Sfta2) in mammals and the effects of Sfta2 knockout in mice. **a,** Schematic diagram showing the correspondence of cell types between mammals and non-mammals, highlighting the emergence of AT1 cells. **b,** UMAP clustering of epithelial cells from four mammalian and non-mammalian species (left). Feature plots showing the expression of four AT1 marker genes across all species (right). **c,** Expression of *sftpb* (AT2 marker gene) and *clic5* (AT1 marker gene) in the lungs of mouse, central bearded dragon, and African lungfish. **d,** Potential protein-protein interaction network based on the String database, showing proteins present in mammalian lung epithelial cells with significantly higher proportions in AT1 compared to AT2 cells. Node size represents the gene’s proportion in mammalian AT1 cells, while color intensity indicates expression level. **e,** Schematic diagram showing the relative positions of sfta2 *and* surrounding genes in African bullfrog, central bearded dragon, mouse, and human. **f,** Expression of *sfta2* across various cell types in the mammalian lungs. **g,** Comparison of respiratory parameters between wild-type (WT, dark bars) and Sfta2 knockout (Sfta2-/-, light bars) mice, showing breathing frequency (FN, breaths/minute), minute volume (mv, total air volume/minute), and tidal volume (v, air volume/breath). **h,** H&E-stained lung sections of Sfta2 knockout homozygous mice compared to wild-type. **i,** Comparison of alveolar wall thickness between *Sfta2* knockout homozygous mice and wild-type, showing significant thickening in knockout mice. **j,** Immunofluorescence images showing F4/80 gene expression in *Sfta2* knockout homozygous mice compared to wild-type.

Notably, the lack of co-expression of *clic5* and *sftpb* in non-mammalian lungs reflect that mammalian AT1 and AT2 cells did not simply differentiate from ancestral lung epithelial cells. Instead, integrating our analysis of chicken and mouse lung epithelial cell developmental trajectories with cell similarity analyses (**Extended Data Fig. 8d**), we can infer that AT2 cells share higher similarity with non-mammalian respiratory epithelial cells. Conversely, AT1 cells appear to be a novel cell type that emerged in mammalian lungs, exhibiting numerous uniquely expressed genes compared to AT2 cells and non-mammalian respiratory epithelial cells (**Extended Data Fig. 8e**). By comparing the expression pattern of AT1 and AT2 cells, we identified 50 protein-protein interactions (PPIs) that have a significant higher proportion in AT1 cells. The potential roles of these PPIs in AT1 cells include caveolae formation and membrane organization (CAV1-CAV2, CAVIN1-CAVIN2)^47^, cytoskeletal support for the thin, extended AT1 morphology (SPTAN1-SPTBN1), and basement membrane formation for proper cell spreading and attachment (LAMC2-LAMA3)^48^. These interactions likely work together to establish and maintain the unique flattened morphology of AT1 cells, which is crucial for their function in gas exchange.

Align with the emergence of new cell type in mammalians, we identified a new gene, *sfta2*, emerged since the common ancestors of mammals (**Fig. 5d and Extended Data Fig. 8f)**. This gene encodes a surfactant protein and is highly expressed in AT2 cells. Consistently, the genes that specifically highly expressed in mammalian epithelial cells are including multiple genes associated with lipid metabolism and synthesis, making the main component of surfactants (**Extended Data Fig. 8g**). Within them, we observed the expression of *klf5*, encoding transcription factor KLF5, in mammalian lungs, but absent in non-mammals. KLF5 regulates surfactant lipid and protein homeostasis, and angiogenic gene expression, influencing lung maturation during the saccular period, when AT1 and AT2 differentiation occurs^49^. These findings are consistent with the unique alveolar structure of mammals, where large curvatures require a rich surfactant system to reduce surface tension^50^. An intriguing question arises regarding the role of *sfta2*, a novel non-surfactant gene, in the context of lung function and evolution (**Fig. 5e**). While most pulmonary surfactant genes, such as those in the Sftp family, are present in non-mammalian species, we sought to investigate whether *sfta2* is essential for maintaining lung function and if it is associated with the emergence of the new AT1 cell type. This inquiry is particularly compelling given that *sfta2* is primarily expressed in AT2 cells and is not consistently highly expressed in AT1 cells across all mammalian species (**Fig. 5f**).

To elucidate the function of *sfta2*, we employed CRISPR/Cas9 technology to generate *sfta2* knockout mice. Our findings revealed that homozygous *sfta2* knockout mice exhibited significantly increased respiratory rates, reduced tidal volumes, and decreased minute ventilation compared to wild-type mice (**Fig. 5g**). Histological examination through H&E staining demonstrated immune cell infiltration in the alveolar walls, resulting in increased alveolar wall thickness (**Fig. 5h-i**). This observation was further corroborated by immunofluorescence imaging of F4/80-positive cells. Furthermore, comparative transcriptomic analysis between *sfta2*^-/-^ and wild-type mouse lungs revealed significant enrichment of upregulated genes in pathways related to wound healing, spreading of epidermal cells, and mitotic cell cycle in *sfta2*^-/-^ mice (**Extended Data Fig. 8h**). Notably, *wnt7a*, which promotes AT2 to AT1 differentiation^51^, showed higher expression in knockout mice. These results collectively indicate that *sfta2* plays an indispensable role in maintaining mouse alveolar cell homeostasis, and may facilitate the emergence of AT1 cells as a new cell type in mammalian lungs.

## Conclusion

Through our exploration of the origins of lungs, we have come to recognize that genetic evidence alone may never definitively answer whether the ancestors of cartilaginous fish possessed lungs. However, contemplating this debate has facilitated a deeper understanding of a more crucial question: what factors ultimately determine the origin of vertebrate lungs. Our systematic analysis underscores the significance of regulatory changes in this evolutionary process. A key finding from our research stems from single-cell analysis of both mature and developing lungs (**Fig. 1** and **Fig. 2**), revealing substantial conservation across species, with hundreds of genes exhibiting shared expression patterns. Lung epithelial cells also display distinctive co-expression patterns (**Fig. 4e**), particularly in genes related to lung surfactants and cell morphology, representing cellular specialization. Notably, these genes are predominantly present in cartilaginous fish, and there is no evidence suggesting a direct correlation between their presence and the existence of lungs. Building upon this foundation, we identified thousands of lung-related CNEs specific to bony fishes. These CNEs are heterogeneously distributed, with high concentrations around genes such as *tbx4* and the *hoxb* gene cluster. Moreover, these CNEs are associated with numerous transcription factors, including *foxa1* and *foxa2*, which exhibit high expression in lung epithelial cells. These findings collectively suggest that the unique regulatory patterns of bony fish, rather than the mere presence of certain genes, are determinative in the presence of lungs.

From an alternative perspective, the role of new genes, especially gene duplicates, in the evolution of lungs cannot be overlooked. Our analysis reveals that approximately one-third of the genes critical for both mature and developing lungs are products of the vertebrate 2R-WGD, a proportion substantially higher than the genome-wide background (**Fig. 3b**). This finding mirrors observations from another study by our team, which demonstrated that the 2R-WGD significantly contributed to the evolution of liver sinusoidal endothelial cells in vertebrates. Furthermore, a new gene shared by mammals, *sfta2*, which was duplicated from the gene *rhcg*, was found to be associated with the novel lung cell type, AT1, in mammals. Knockout experiments demonstrated that the absence of *sfta2* results in significant respiratory deficiencies in mice, increased alveolar wall thickness, and notable inflammation. Although *sfta2* is not directly expressed in AT1 cells but rather in AT2 cells, these results indicate that its expression may play a crucial protective role essential for the existence of AT1 cells. These observations highlight the significant impact of gene duplication events on the origin and specialization of lung structures in vertebrates.

The evolution of lungs also corroborates two previous speculations about the evolution of complex organs. Firstly, it supports the idea that the evolution of lungs was a gradual process, consistent with Darwin’s predictions^52^. Similar to the eye and other organs^6, 53, 54^, lung evolution appears to have occurred incrementally over time. While epithelial cells have an ancient origin, the respiratory epithelial cells of bony fishes exhibit unique expression patterns and even novel core regulatory complexes. The evolutionary progression of respiratory epithelial cells demonstrates a gradual transition from aquatic to terrestrial environments, followed by additional specialization in mammals. Secondly, our findings echo Jacob’s assertion that evolution acts as a “tinkerer”^55^, using existing genetic foundations, including the recruitment and co-option of pre-existing genes and regulatory elements ^4, 56, 57^. Overall, these results imply that modifications to regulatory networks are crucial in the origin and evolution of this organ and that the emergence of new genes or gene duplicates provides essential materials, even if these genes do not immediately come into play.

## Methods

### Ethics statement and processing of samples

The experimental animals used in this study include adult white spotted bamboo shark, Senegal bichir, African lungfish, African bullfrog, central bearded dragon and chicken embryos. Chicken embryos were purchased from farms in Yangling, Shaanxi Province, China and other animal samples were purchased from pet markets in Guangzhou and then identified in School of Ecology and Environment, Northwestern Polytechnic University. All experimental protocols and animal use for this study were approved by the Northwestern Polytechnic University Ethics Committee Institutional Review Board (202301024 and 202301143).

To generate scRNA-seq data for adult Senegal bichir and African lungfish, after anesthesia (using Ms-222 100 mg/l,) and exposure of the body cavity, a 0.5 cm incision was made in the liver. Perfusion with PBS was then conducted through the heart to flush out as much blood as possible before tissue sampling. Tissue samples were cut into small pieces in ice-cold PBS and then digested with corresponding enzymes to prepare single cell suspensions. Single cell RNA sequencing was conducted on lung using a Chromium Controller (10× Genomics, US), while the other tissues using DG1000 (BioMarker Technologies, Chain).

For the terrestrial adult animals, including the African bullfrog, central bearded dragon, and chicken, the same tissue sampling and single-cell suspension preparation methods were used, except that the animals were anesthetized using a ketamine hydrochloride injection (10 mg/l) instead of MS-222. For adult white spotted bamboo shark, The suspensions of each tissues were then subjected to single cell RNA sequencing using C4 (BGI, Chain).The lungs/swim bladder of adult African bullfrog, central bearded dragon and zebrafish were sampled for single cell RNA sequencing with 10× Genomics.

We additionally collected chicken lung samples at embryonic stages on days 6, 7, and from postnatal day 3. Embryonic lung samples, obtained under a dissecting microscope, were pooled (30 lungs per pool) to form a single sample. Postnatal day 3 lung samples were collected using the same methodology as employed for other adult lung samples. Subsequently, all chicken lung samples underwent single-cell RNA sequencing using 10× Genomics. Among all samples, there were 2 biological replicates in the lungs of Senegal bichir, African lungfish and African bullfrog and 2-3 biological replicates in the tissues of the white-spotted bamboo shark. Details of all samples including biological replicates and quality are listed in Supplementary Table 1.

### Genome sequencing, assembly and annotation

Fresh muscle tissue samples from adult central bearded dragon were collected and subjected to high-throughput sequencing for genome assembly. Genomic DNA was extracted from the samples using a DNA purification kits (Qiagen, 69516) and sequenced on Illumina HiSeq2000 and Oxford Nanopore GridIOn platforms (ONT), generating 140 Gb and 190 Gb of data, respectively. For Hi-C sequencing library construction, 185 Gb of data was obtained from the Illumina HiSeq X Ten platform.

We estimated the genome size to be 1.8 Gb based on k-mer frequencies using Jellyfish (version 2.2.10)^58^ and GenomeScope^59^. To assemble contig-level genomes, NextDenovo software (version 2.4.0; https://github.com/Nextomics/NextDenovo) was used to self-correct the long reads from ONT platform and resulting contigs were then polished using NextPolish (version 1.3.0)^60^. The Hi-C short reads were aligned to the contigs using Juicer (version 1.6)^61^, and the contigs were anchored using 3D-DNA (version 180419)^62^. The Juicebox Assembly Tools (https://github.com/theaidenlab/juicebox). were applied to manually correct the connect. Finally, the quality of the genome assembly, 93% completeness was assessed using BUSCO (version 4.1.2)^63^ with the tetrapod lineage database.

To annotate the protein-coding genes in genome, we generated *ab initio* predictions using Augustus (version 3.2.1)^64^ with internal gene models and homolog-based annotation based on the protein of eight species including human (*Homo sapiens*; GRCh38), mouse (*Mus musculus*, GRCm38), chicken (*Gallus gallus*, GRCg6a), green anole (*Anolis carolinensis,* ensemble GCA_000090745.2 release), Indian cobra (*Naja naja;* ensembl GCA_009733165.1 release), Chinese softshell turtle (*Pelodiscus sinensis;* NCBI GCF_000230535.1 release), Pinta island tortoise (*Chelonoidis abingdonii,* ensembl GCA_003597395.1 release), common snapping turtle (*Chelydra serpentina*; ensembl GCA_007922165.1 release) and protein sequences of transcripts of central bearded dragon. These proteins, only the longest transcript from each gene were aligned to the genome using the default parameters of BLAT (version 35.1)^65^. Finally, gene prediction performed by GeneWise (version 2.4.1)^66^ and manually removed the duplicate annotations. 83% completeness of the gene set was assessed using BUSCO^63^ with the tetrapod lineage database.

### Processing of scRNA-seq

To ensure consistent gene nomenclature across multiple species included in this study, gene names for each species were converted to human gene names based on sequence homology. The Reciprocal Best Hits (RBH) method for BLASTP (version 2.9.0)^67^ was utilized to identify one-to-one homologous genes between human and each species. Genes without a one-to-one homologous match were retained with their original species-specific names.

To generate expression matrix, in addition to the central bearded dragon genome generated in this study (PRJNA1026724 in NCBI), genomes of white spotted bamboo shark genome (GCF_004010195.1), Senegal bichir (GCA_016984135.1), African lungfish (GCA_019279795.1), African bullfrog (GCA_004786255.1) and chicken (GRCg7b) were obtained from NCBI. The genome annotation of African bullfrogs was obtained using the above method in the same way as that of central bearded dragon, and the annotations of other genomes were also obtained from NCBI. Single cell RNA raw reads from 10× Genomics, DG1000, C4 sequencing platforms were aligned to corresponding genomes with default parameters using Cell Ranger (https://github.com/10XGenomics/cellranger), BSCMatrix (http://www.bmkmanu.com/portfolio/tools), DNBC4tools (https://github.com/MGI-tech-bioinformatics/DNBelab_C_Series_HT_scRNA-analysis-software) respectively and obtained expression matrices.

The expression matrices of mammalian lungs were obtained from GSE133747^27^. For each sample, we used the Read10X and CreateSeuratObject functions from the Seurat (version 3.1.1)^68^ R package to generate a Seurat object from the expression matrix and use the parameters min.cells=3, min.features= 200 to filter out genes expressed by less than 3 cells and cells expressed by less than 200 genes. Scrublet^69^ and SoupX (version 1.5.0)^70^ were employed to remove doublets cells and ambient RNA with default parameters. Integration of biological replicates for tissues of the white spotted bamboo shark and lungs of adult chicken and mammalian lungs was performed by merge in R and we checked no batch effect among replicates. Data normalization was conducted using the NormalizeData function, and the top 2,000 most variable genes were identified with the FindVariableFeatures function. Subsequently, the data was scaled using the ScaleData function, and principal component analysis (PCA) was performed for dimensional reduction using the RunPCA function. Cell clustering was carried out using the FindNeighbors and FindClusters functions, followed by dimensionality reduction mapping of cell clusters using RunUMAP. Given that the merge method did not sufficiently eliminate batch effects for two biological replicates of lungfish and frog lungs, an additional step was introduced post-PCA: employing the RunHarmony function from the harmony package for integration. After obtaining cell clusters marker gene identification for each cluster was performed using the FindAllMarkers function with default parameters. Erythrocytes were excluded based on marker genes (*hba2*, *hbb-bh1*, *alas2*) across all species’ lung tissues. In summary, the lungs of Senegal bichir and African lungfish, African bullfrog, central bearded dragon, chicken, pig, human, mouse, and rat obtained 12,034, 13,925, 6,056, 6,468, 47,003, 11,166, 12,677, 7,744, and 10,787 effective cell numbers respectively. The effective cell numbers of other tissues from white spotted bamboo shark, Senegal bichir and African lungfish as well as lungs from chicken at different developmental stages, and all data quality control results including the cell number and gene expression number of each cell type are shown in the in Supplementary Table 1. In addition, cell annotations of non-mammalian lungs and the expression of some marker genes in are provided in **Extended Data Fig. 1.**

### Integration of scRNA-seq of lung across different species

To integrate 127,860 lung cells across different species, we employed the SAMAP tool (version 1.0.13)^71^ in python3. First, we converted cleaned scRNA-seq data from Seurat objects into h5ad format using the SaveH5Seurat and Convert functions within the SeuratDisk package in R. Next, we employedsmap_genes.sh’ to conduct pairwise comparisons of proteins among 9 species, ensuring the identification of orthologous genes across all species. Subsequently, we utilized the SAMAP function to construct objects and the sm.run function for data integration, with parameters set as ‘pairwise=True, N_GENE_CHUNKS=4, crossK=5’. We then employed the scanpy package to annotate cells based on the SAMAP integration results. Cell clustering was performed using the neighbors function with parameter n_neighbors=50, and the leiden function with parameter resolution=0.5. Cell clusters were annotated based on human and mouse marker genes. For each cell type, if there are fewer than 30 cells of that cell type in a given species, it is considered that the group does not exist in that species.

To further ensure the accuracy of cross-species integration, we implemented multiple methods based on the expression of 6,303 orthologous genes: 1) Canonical Correlation Analysis (CCA): We utilized the FindIntegrationAnchors function with dims=1:30 and IntegrateData functions within Seurat, followed by normalization using ScaleData. RunUMAP was used to obtain the dimensionality reduction matrix of the cells. 2) Reciprocal Principal Component Analysis (RPCA): We employed the FindIntegrationAnchors function with parameters ‘reduction=”rpca”, k.anchor=80’, followed by integration using the IntegrateData function with dims=1:30. UMAP matrix was obtained using the same method as CCA. 3) Mutual nearest neighbors (MNNs)^72^: Integration was carried out using the RunFastMNN function within the SeuratWrappers package with default parameters. Dimensionality reduction was obtained using the RunUMAP function with reduction=”mnn”. 4) LIGER (linked inference of genomic experimental relationships)^73^: We merged lung data from each species using the merge function, followed by data standardization through NormalizeData and identification of variable genes using FindVariableFeatures. After scaling data using ScaleData with parameter do.center = F, integration was then performed using the RunOptimizeALS function in the rliger package with parameters k=30, lambda=5, and RunQuantileNorm function. Cell dimensionality reduction was obtained through RunUMAP. 5) scVI and scANVI^74^: We employed the scVI and scANVI probabilistic models from scvi-tools to map cell atlas manifolds across different species. The tuning of hyperparameters for these models was conducted using the Ray Python library (version 2.11.0). The final parameters were set as follows: n_hidden=256, dropout_rate=0.25, learning rate (lr)=5e-4, and max_epochs=1000. We mapped cell annotations based on SAMAP to UMAP plots generated from the integration results of the six methods. UMAP plots of CCA, RPCA, and MNN demonstrated consistent cell clustering, affirming the accuracy of cell integration and annotation (see **Extended Data Fig. 2c**)

### Processing of bulk RNA-seq

For transcriptome analysis, we collected RNA samples from 18 tissues (ampullae, bone, brain, esophagus, liver, fin, gill, heart, inner ear, intestine, kidney, lateral line, muscle, pancreas, skin, spleen, stomach, tooth) with 2-3 biological replicates for each tissue of white spotted bamboo sharks. Tissues were snap-frozen in liquid nitrogen, and then total RNA for tissues was extracted using the mirVana™ miRNA isolation kit (Ambion-1561, Thermo Fisher Scientific, Inc, USA). RNA purity and integrity were detected using a NanoPhotometer spectrophotometer and an Agilent 2100 bioanalyzer (Agilent Technologies, CA, USA). Library construction was performed using Illumina NEBNext® UltraTM RNA Library Prep Kit (Illumina, Inc, USA) and sequencing on the Illumina NovaSeq 6000 platform to generate 150 bp paired-end reads. The process is completed by China’s Tianjin Novotech Technology Co., Ltd. Clean reads were obtained raw data using Fastp (version 0.20)^75^ and were mapped to the white spotted bamboo shark genome (NCBI GCF_004010195.1 release) using HISAT2 (version 2.2.0)^76^. While in this alignment process, environmental RNA will not be aligned to the target genome. FPKM values of each sample were calculated using StringTie (version 2.1.3b)^77^ with -eB options. Raw RNA-seq data of brain, gills, heart, liver, muscle and skin for Senegal bichir and African lungfish were obtained from PRJNA599026 at NCBI and from PRJCA002950 at the National Genomic Data Centre (https://bigd.big.ac.cn/bioproject/) respectively. The reference genomes used were GCA_016984135.1 (Senegal bichir) and GCA_019279795.1 (African lungfish). FPKM values were obtained using the same method as above.

To analyze the comparison of gene expression for multiple tissues between Senegal bichir, African lungfish and white spotted bamboo shark, we used the ComBat function in R to adjust for species batch effects and then used the prcomp function in the ape package (version 5.7.1) in R to complete the PCA analysis.

### Processing of CUT&Tag

We collected chicken lung samples at embryonic stages on days 9, 11, and 19, and rapidly froze them in liquid nitrogen immediately after sampling. Cells were harvested via the Nuclear Extraction Kit (SN0020, Solarbio, Beijing, China) for CUT&Tag assay using Hyperactive Universa CUT&Tag Assay Kit (TD903-02, Vazyme, Nanjing, China). Cells were bound to concanavalin A-coated magnetic beads and permeabilized using the non-ionic decontaminant digitalis saponin. The transposon fused to Protein A/G is precisely targeted to cleave DNA sequences in the vicinity of the target protein mediated by a primary antibody against the target protein, including H3K4me1 (Abcam, USA) and H3K27ac (Abcam, USA), the secondary antibody and Protein A/G. The transposon is cleaved with a splice sequence at both ends of the cleaved fragment, which can be amplified by PCR to form a library for genomic sequencing using the Illumina NovaSeq 6000 platform. We conducted 2 replicates for each marker. For sequencing data analysis, the chicken genome (GRCg6a) index was constructed and the reads were aligned to the reference genome using bowtie2 (version 2.4.4)^78^ with the parameters “--end-to-end --very-sensitive --no-discordant --phred33 -I 10 -X 700” after filtering out low-quality sequences and removing adapters using fastp^75^. The resulting alignment files (SAM) were sorted, and various file format conversions were performed using SAMtools (version 1.9-74-gf69e678)^79^, BEDtools (version 2.29.2)^80^, and DeepTools (version 2.0)^81^. MACS3(version 3.0.0b3)^82^ was used with the options “-q 0.05 -- keep-dup all --nomodel” after removing redundant reads with Picard (http://broadinstitute.github.io/picard/). Putative enhancers were defined when H3K4me1 and H3K27ac peaks overlapped.

### Identification of highly expressed genes, lung specific genes, TF regulons and cell communication pathways in lungs

First, we categorized the lung cells across 9 species used in this study into 4 major populations (epithelial cells, endothelial cells, stromal cells, immune cells). Subsequently, we employed the Wilcoxon test using the FindMarkers function in Seurat version 4 to identify DEGs for each cell population in each species with parameters min.pct=0.05, logfc.threshold=0.1. and adjusted p-values ≤ 0.05. Given the species-specific gene expression patterns, we selected DEGs present in at least six out of the nine species, including both lungfish and bichir, for each cell population. These DEGs were considered as conserved highly expressed genes for the respective cell populations in vertebrate lung tissues. We obtained a total of 401 DEGs shared by lung tissues, including 69 genes for epithelial cells, 90 genes for endothelial cells, 168 genes for stromal cells, and 74 genes for immune cells.

To identify genes specifically expressed in lung tissue rather than those intrinsic to cell populations, we obtained scRNA-seq data from 10 tissues in lungfish and 9 tissues in bichir. Cells were classified into the four aforementioned populations, and the average expression of each cell type was calculated. The expression of each conserved lung gene. To identify genes specifically expressed in lung tissue rather than those intrinsic to cell populations, we obtained scRNA-seq data from 10 tissues in lungfish and 9 tissues in bichir. Cells were classified into the four aforementioned populations, and the average expression of conserved highly expressed genes for each cell population was calculated.

We used a two-step process to identify lung-specific genes. First, for each species, the expression of each lung conserved highly expressed gene was scaled across all cell populations, and a one-tailed Z test was used to determine whether their expression in a specific lung cell type was significantly higher than in other cells (p value < 0.05). Second, for the genes identified in the first step, such as those obtained in epithelial cells, their expression was scaled across epithelial cells of all tissues. The same method was then used to select genes with lung-specific expression. Genes identified simultaneously in both species were considered conserved lung-specific genes. As a result, we identified 11 lung-specific genes in epithelial cells, 7 in endothelial cells, and 13 in stromal cells, but none in immune cells.

To infer the activated gene regulatory network (TF regulon) in the lung of each species, we utilized the SCENIC (version 1.2.4)^83^ packages in R, leveraging human TF (transcription factor) data as a reference. Regulons active across all species were defined as conserved TF regulons. We used CellChat (version 1.0.0)^84^ in R to infer cell-cell communication pathways in the lung of each species and VEGF signaling networks identified in all species were considered as conserved cell communication pathway.

### Identification of key genes for lung development

We downloaded published single-cell transcriptome data of the mouse embryonic lung, including data from embryonic days 12, 15, and 18, and postnatal days 0, 3, 5, 7, and 14. The gene expression matrices were obtained and cells were clustered and annotated according to marker genes using the same method as above for each sample. We selected epithelial cells from each sample, merged them, and used the Seurat package to perform dimensionality reduction on the epithelial cells based on the top 2000 highly variable genes to obtain UMAP matrices. We then used the slingshot package (version 2.2.1)^33^ package in R to infer the cell trajectories. This pseudotime results was confirmed using the R package Monocle (version 2.26.0)^85^. To identify genes with significant expression variation along the trajectory, we applied the differentialGeneTest function from Monocle in R, identifying genes with a q-value < 1e-6. In mice, we identified three trajectories. Using the same method, we constructed a trajectory for epithelial cells in the chicken lung across four stages and identified highly variable genes using same method. We found 270 genes that were gradually highly expressed both in chicken and mouse pseudo-time trajectories.

We compared the similarity in gene expression of cells along the trajectories between chicken and mouse. To achieve this, we divided the cells along the mouse trajectories into different stages. For the AT2 cell trajectory in mice, we divided it into six stages and for chicken trajectories, we divided them into four stages. We then calculated the average expression of 1:1 orthologous genes between chicken and mouse for each stage. Subsequently, we computed the correlation between the different stages of chicken and mouse using the cor function in R.

To identify genes highly expressed in stromal cells, taking mouse as an example, we identified the top 3000 highly variable genes (considering only the 1:1 orthologous genes between chicken and mouse) in the mouse lung development scRNA-seq data. Among these 3000 genes, we identified those highly expressed in stromal cells using the FindMarkers function with the parameters logfc.threshold = 0.15 and min.diff.pct = 0.1. The same method was applied to identify genes highly expressed in chicken stromal cells. The genes shared between chicken and mouse were considered as the final gene set critical for lung development.

### Origin of lung specific genes

To infer the origin of lung-related genes, including highly expressed conserved lung genes and key genes involved in lung development, we performed the following steps. Due to the large number of genes, we first downloaded the protein sequences of 14 cartilaginous fish species, including white spotted bamboo shark,brownbanded bamboo shark (*Chiloscyllium punctatum*; NCBI GCA_003427335.1 release), great white shark (*Carcharodon carcharias*; NCBI GCF_017639515.1 release), elephant shark (*Callorhinchus milii*; NCBI GCF_000165045.2 release), Epaulette shark (*Hemiscyllium ocellatum*; NCBI GCA_020745735.1 release); Cloudy catshark (*Scyliorhinus torazame*; NCBI GCA_003427355.1 release); whale shark (*Rhincodon typus*; NCBI GCF_021869965.1 release); Little skate (*Leucoraja erinacea*; NCBI GCA_028641065.1 release); Small-eyed rabbitfish (*Hydrolagus affinis*; NCBI GCA_028641065.1 release); Spiny dogfish (*Squalus acanthias*; NCBI GCA_030390025.1 release); Zebra shark (*Stegostoma tigrinum*; NCBI GCA_030390025.1 release); thorny skate (*Amblyraja radiata*; NCBI GCF_010909765.2 release), Small-spotted catshark (*Scyliorhinus canicula*; NCBI GCA_902713615.2 release) and Smalltooth sawfish (*Pristis pectinata*; NCBI GCA_009764475.2 release). We conducted a reciprocal best hits (RBH) analysis between all mouse proteins and each cartilaginous fish proteomes using the same method as described above. If a target gene was matched in more than two cartilaginous fish species, we considered that these genes present in cartilaginous fish. For the remaining genes, we selected the proteomes of 24 deuterostome species, including roundworms (*Caenorhabditis elegans*; NCBI GCF_000002985.6 release), sea urchin (*Strongylocentrotus purpuratus*; NCBI GCA_000002235.5 release), starfish (*Asterias rubens*; NCBI GCF_902459465.3 release), lancelet (*Branchiostoma floridae*; NCBI GCF_000003815.2 release), sea squirts (*Ciona intestinalis*; NCBI GCF_000224145.2 release), hagfish (*Eptatretus burger*; NCBI GCF_024346535.1 release), lamprey (*Petromyzon marinus*; NCBI GCF_010993605.1 release), elephant shark, thorny skate, great white shark, white spotted bamboo shark, Senegal bichir, spotted gar (*Lepisosteus oculatus*; ensembl GCA_000242695.1 release), European eel (*Anguilla anguilla*; NCBI GCF_013347855.1 release), stickleback (*Gasterosteus aculeatus*; ensembl BROAD S1 release), large spiny eel (*Mastacembelus armatus*; ensemble GCA_900324485.2 release), zebrafish (GRCz11), African lungfish, xenopus (*Xenopus tropicalis*; ensembl 9.1 release), African bullfrog, central bearded dragon, chicken, human and mouse. For each gene, the human protein sequence was aligned to the proteome of each selected species using Diamond (version 2.1.4.158)^86^ with the parameter --ultra-sensitive. Hits with *P*-value > 1e^-5^ were filtered out and if more than 5 hits were aligned, only the top 5 hits were retained for subsequent analysis. These proteins were aligned using MAFFT (version 7.407)^87^ and phylogenetic trees were constructed using RAxML (version 8.2.12)^88^ with the parameter “-f a -m PROTGAMMAAUTO -T 50 -N 100” for each gene. When sequences from a given lineage had paralogs not present in outgroup species on the phylogenetic tree, the gene was considered a new duplicate in that lineage.

### Selected pressure detection for lung specific genes

To assess selection pressure for lung specific genes, we calculated the Ka and Ks for these genes. For each gene, the protein sequences of the 1:1 orthologs of the corresponding species were first identified in the phylogenetic gene tree. The coding sequences (CDS) of the orthologs were aligned utilizing the prank (version 170703)^89^. To refine the alignment and remove gaps, we employed Gblocks (version 0.91b)^90^ to trim gapped CDS regions. The codeml of Paml (version 4.9j)^91^ were used to reconstruct the ancestral sequences. For each gene, the Ka and Ks were calculated by KaKs_Calculator (version 3.0) comparing ancestral sequences and no gap CDS sequences for each species.

### Origin of lung-related conserved noncoding elements

To identify CNEs within H3K27ac peaks in chicken and mouse, we first selected 23 amniotes genomes including central bearded dragon, common wall lizard (*Podarcis muralis*; ensembl GCA_004329235.1 release), spectacled caiman(*Caiman crocodilus*; NCBI GCA_030014935.1 release), Abingdon island giant tortoise (*Chelonoidis abingdonii*; NCBI GCF_003597395.1 release), burmese python (*Python bivittatus*; NCBI GCF_000186305.1 release), corn snake (*Pantherophis guttatus*; NCBI GCF_001185365.1 release), western painted turtle (*Chrysemys picta bellii*; NCBI GCF_000241765.5 release), red eared slider (*Trachemys scripta elegans*; NCBI GCF_013100865.1 release), turkey (*Meleagris gallopavo*; ensembl GCA_000146605.4 release), new caledonian crow (C*orvus moneduloides*; NCBI GCF_009650955.1 release), zebra finch (*Taeniopygia guttata*; ensembl GCA_003957565.2 release), platypus (*Ornithorhynchus anatinus*; ensembl GCA_004115215.2 release), tammar wallaby (*Notamacropus eugenii*; ensembl GCA_000004035.1 release), dog (*Canis familiaris*; ensembl GCA_014441545.1 release), African savanna elephant (*Loxodonta Africana*; ensembl GCA_000001905.1 release), Hoffmann’s two-fingered sloth (*Choloepus hoffmanni*; NCBI GCA_000164785.2 release), duck(*Anas platyrhynchos*; NCBI GCF_015476345.1 release), Chinese alligator (*Alligator sinensis*; NCBI GCF_000455745.1 release), cattle (*Bos taurus;* NCBI GCF_002263795.1 release), gray short-tailed opossum (*Monodelphis domestica*; NCBI GCF_000002295.2 release), chicken, human and mouse to identify CNEs. The whole genome alignments of all species were generated by LAST (version 1282)^92^ with parameters “-uNEAR” when index the reference genome, using the genome of mouse as a reference. Multiz (version 11.2)^93^ was subjected to obtain multiple genome alignments and then PhastCons (version 1.6)^94^ was used to get conserved elements with parameters “--target-coverage 0.3 --expected-length 45 --rho 0.3”. We filtered all CDSs overlapped with them to obtain conserved non-coding elements and filter CNEs less than 50 bp. Then, we obtained the H3K27ac ChIP-seq peak bed files of mouse embryonic lungs from the ENCODE database and the H3K27ac CUT&Tag peak bed files of chicken embryonic lungs from this study. We identified CNEs located within these H3K27ac peaks and traced the origins of these CNEs.

To explore the origin of these CNEs, we used BLASTN (version 2.9.0)^67^ to align these CNEs to lancelet, hagfish, lamprey, cloudy catshark, thorny skate, raja gigas (*Pristis pectinate*, NCBI GCF_009764475.1 release), great white shark, Whitespotted bamboo shark, brownbanded bamboo shark, whale shark, Senegal bichir, spotted gar, alligator gar, bowfin (*Amia calva;* NCBI GCA_017591415.1 release), American paddlefish (*Polyodon spathula;* NCBI GCF_017654505.1 release), common toad (*Bufo bufo*; NCBI GCF_905171765.1 release), microcaecilia (*Microcaecilia unicolor*; NCBI GCF_901765095.1 release), European eel, coelacanth (*Latimeria chalumnae*; ensembl GCA_000225785.1 release), xenopus, two-lined caecilian (*Rhinatrema bivittatum*; NCBI GCF_901001135.1 release),African bullfrog with the parameters “-word_size 10 -evalue 1e-5 -reward 1 -penalty-1 -gapopen 2 -gapextend 2 -dust no -soft_masking false -culling_limit 20”.The origin of the CNEs was inferred based on their alignment results.

To obtain the potential linear relationships between genes and CNEs, we used miniprot (version 0.12)^95^ to align the mouse protein sequences of lung-conserved genes and key genes for lung developmental in to the 41 high-quality genomes mentioned above. For each species, we recorded the genomic positions of the genes based on miniport result. Simultaneously, based on the previously mentioned CNE alignment results, we obtained the positions of CNEs within a 2Mb region adjacent to each gene. In this way, we obtained CNE and gene pairs that may interact. To further validate the interactions between CNEs and genes, we utilized lung data from the 3D Genome Browser (http://3dgenome.fsm.northwestern.edu/) to analyze the spatial interactions. The alignments of CNEs were manually checked and plotted using VISTA (version 1.4.26)^96^.

TFBS searching on CNEs were conducted with fimo from MEME Suite (version 5.4.1)^97^. PWMs (position weight matrix) were obtained from JASPAR (2023)^98^ and HOCOMOCO (version 11)^99^ database comprehensively. Each CNE from candidate species were scanned respectively, only those motifs simultaneously discovered in all species were defined as a true TFBS signal.

### Identification of gene pairs or protein-protein interaction (PPI) in new cell type

To identify gene pairs in lung respiratory epithelial cells using single-cell RNA sequencing data, we first selected 69 genes highly expressed in epithelial cells and randomly generated gene pairs and then calculated the co-expression ratios of gene pairs in lung epithelial cells from nine vertebrate species and in 55 different cell types from bamboo shark. For each gene pair, the co-expression ratio was defined as the proportion of cells expressing both genes within a given cell type. We then compared the co-expression ratios in lung epithelial cells against the highest co-expression ratio found among the shark cell types. Gene pairs were considered significant if they met two criteria: (1) their co-expression ratio in lung epithelial cells showed a z-score < -1.65 when compared to the distribution of ratios in shark cell types, and (2) their co-expression ratio in lung epithelial cells was more than 5-fold higher than the maximum ratio observed in any shark cell type.

To identify the protein-protein interaction (PPI) networks in AT1, we utilized the STRING protein interaction database to identify interaction pairs among conserved highly expressed genes in AT1 cells across four mammalian species. We filtered out interactions identified solely through text mining to retain high-confidence interaction pairs. For each PPI pair, we calculated the proportion of cells expressing both genes in AT1 and AT2 of each species. We then performed a one-tailed paired t-test in R to identify PPIs with a significantly higher proportion in AT1 cells (p.adjust < 0.05). and identified PPIs with a high proportion of interactions in AT1 cell types compared with AT2 cell types.

### proteome analysis of gills and lungs

In order to detect the protein abundance of SFTPC in lungfish gills and lungs, lungfish were dissected to obtain gill and lung tissue samples, with two biological replicates for each tissue type. Samples were taken from -80°C and ground to a fine powder in liquid nitrogen. Each sample was mixed with four volumes of lysis buffer (1% SDS, 1% protease inhibitor) and sonicated. After centrifugation at 12,000 g for 10 minutes at 4°C, the supernatant was collected, and protein concentration was determined using a BCA kit. Then, Equal amounts of protein from each sample were adjusted to the same volume with lysis buffer. Pre-chilled acetone (1 volume) was added, vortexed, followed by four volumes of pre-chilled acetone, and precipitated at -20°C for 2 hours. The precipitate was washed with cold acetone, dried, and resuspended in 200 mM TEAB. Trypsin was added at a 1:50 ratio (enzyme:protein) and digested overnight. Samples were reduced with 5 mM DTT at 56°C for 30 minutes and alkylated with 11 mM IAA at room temperature for 15 minutes in the dark to obtain the final peptide fragment. The peptides were dissolved in liquid chromatography mobile phase A and separated using a Vanquish Neo ultra-high performance liquid chromatography system to generate raw data for mass spectrometry detection. Proteins were identified using the lungfish genome as the database with the DIA-NN (version 1.8)^100^ search engine under default parameters. The enzyme mode was set to Trypsin/P with a maximum of one missed cleavage. The fixed modifications were set to: N-term M excision and C carbamidomethylation. The search results need further data filtering, and the filtering conditions were set as follows: Precursor and protein FDR are set to 1% and the identified protein must contain at least one unique peptide segment. Normalized intensity (I) is transformed by centering to obtain the relative quantitative value (R) of the protein in different samples. Rij=Iij/Mean(Ij). We focus the relative value of the three copies of SFTPC in the lungs and gills.

### Highly expressed genes for lobe-fined, tetrapod and mammalian lung

To identify lobe-fined fish originated highly expressed genes in given cell population, taking epithelial cells as an example (significantly higher expressed in epithelial cells compared with other cells, see method aforementioned), genes highly expressed in the epithelial cells among all lobe-fined fishes (One species other than lungfish is allowed to escape), but not in the Senegal bichir, were considered as lobe-fined fish highly expressed genes in epithelial cells. Genes highly expressed in the epithelial cells among all six terrestrial species (One species other than African bullfrog is allowed to escape) but not in epithelial cells from both the Senegal bichir and lungfish were considered terrestrial species specific highly expressed genes. Similarly, for mammals, genes with high expression in epithelial cells in all mammalian species without high expression in the epithelial cells of any of the remaining individual species were considered as mammalian highly expressed genes in epithelial cells. Functional enrichment of highly expressed genes was achieved by the clusterProfiler package^101^ in R. The same method was applied to define genes highly expressed in endothelial cells and stromal cells.

### *In situ* hybridization

Lungs and other tissues from Senegal bichir, African lungfish, African bullfrog, central bearded dragon and mouse were sampled and fixed in situ hybridization fixation solution at room temperature for 24 hours. After dehydration with gradient alcohol, the tissue was embedded in paraffin. Paraffin slices were dewaxed with xylene, boiled in the repair liquid for 10 minutes and naturally cooled. The sample in slices was circled with a Pap Pen. Proteinase K (20ug/ml) was digested at 37°C for 15 minutes. After washing with pure water, PBS was used in washing 3 times for 5 minutes each time. Samples were treated with 3% methanol-H_2_O_2_ and incubated at room temperature in the dark for 15 minutes. The slides were placed in PBS (pH7.4) and washed on a shaker 3 times for 5 minutes each time. A total of eight probes were designed for hybridization, and probe information is provided in the supplementary table 10. For each probe, prehybridization solution was dripped in slides and incubated at 37°C for 1h. The prehybridization solution was removed and the probe hybridization solution was dripped at a concentration of 500nM. Hybridization was performed overnight in a thermostatic incubator at 42°C. After removing the hybridization solution, Imaging oligo (DIG) hybridization solution at 42°C was incubated for 3h. Then 2×SSC, 37°C washed for 10min, 1×SSC, 37°C washed 2×5min, 0.5×SSC 37°C washed for 10min. Rabbit serum was dripped at room temperature for 30min, the blocking solution was poured, and anti-DIG-HRP was dripped. Incubate at 37°C for 50 min, then wash with PBS for 4 times, 5 min each time. FITC-Tyramide reagent was dripped and reacted at room temperature in the dark for 5min. Then TBST was washed 3×10min, and PBS was washed 5min. DAPI was dripped onto the sections, incubated in the dark for 8min, rinsed and then mounting. The sections were observed, and images were acquired on an inverted fluorescence microscope (NIKON Eclipse ci, JAPAN).

### CRISPR gene editing for *Sfta2* mouse and functional assessment

To establish Sfta2 knockout mice, the following steps were undertaken:1, Cas9 mRNA and gRNA were synthesized through in vitro transcription; 2, The mixture of Cas9 mRNA and gRNA was microinjected into fertilized eggs of C57BL/6J mice; 3, The injected embryos were implanted into surrogate mothers to produce F0 mice, which were then identified via PCR and sequencing; F0 positive mice were crossed with wild-type C57BL/6J mice to obtain 20 F1 offspring. The genotypes of F1 mice were confirmed by PCR and sequencing. These steps were conducted by the company of Shanghai Model Organisms.

We utilized whole-body plethysmography (WBP; TOW TECH, China) to assess the lung function of 4-week-old homozygous and wild-type mice (three male mice each). The parameters measured included respiratory frequency (F: number of breaths per minute), tidal volume (VT: amount of gas inhaled or exhaled during each breath.), and minute volume (MV: minute ventilation). We first ensure the WBP equipment is airtight and properly calibrated before use. Mice were placed individually in the sealed chamber and after the mice adapt for 20 minutes, start recording the data for 10 minutes. Take the average value of the observation value every 2 minutes. Each indicator of each mouse corresponds to 5 values. The differences between the experimental and control groups were analyzed using the t.test function in R to determine statistical significance.

To measure alveolar septum thickness, we first obtained 100× magnification images were taken of random alveolar regions using CaseViewer 2.4 scanning software based on results of hematoxylin-eosin staining. Image-Pro Plus 6.0 analysis software was used to measure the thickness of any 5 alveolar septa in each slice with millimeters as the standard unit. The t.test function in R was used to test whether the difference between the experimental group and the control group was significant.

To observe macrophages in the lung, we performed the following procedure on lung tissue sections from both control and experimental groups. After deparaffinization, we used absorbent paper to remove excess liquid around the tissue on the slides. We then used an immunohistochemistry pen to draw a circle around the tissue. Next, we added 3% BSA to the tissue to block nonspecific binding and incubated for 30 minutes. We added the primary antibody (F4/80, diluted 1:1000) and incubated the sections at 4°C overnight. The sections were then washed three times with PBS solution, each wash lasting 5 minutes. We added the fluorescent secondary antibody (CY3-conjugated goat anti-rabbit IgG, diluted 1:300) and incubated in the dark at room temperature for 50 minutes. After another three washes with PBS solution (5 minutes each), we added DAPI staining solution and incubated in the dark at room temperature for 10 minutes. Finally, the sections were mounted with an anti-fade mounting medium. The tissue sections were observed using an upright laser scanning confocal microscope.

### Dual-luciferase assay

The chemically synthesized fragments of mouse CNE1 (GRCm38: 11:85428174-85428564), These CNE fragments were inserted into the pGL4-promoter vector (Promega, USA) and sequenced (TsingKe Biotech, China). Cells were seeded in 24-well plates and after 24 hours, 400 ng of the constructed pGL3-promoter-CNE vector and 40ng pRL-TK vector were co-transfected into A549 cells using PEI reagent (yeasen, China). After 36 hours of transfection, cell lysates were collected using Passive Lysis Buffer. Luciferase activity was measured by Multi-Mode Microplate Reader (BioTek, USA) using a dual luciferase reporter system kit (Promega). Dual fluorescence reporter results were obtained by dividing Firefly Luciferase activity by Renilla luciferase activity.

## Acknowledgments

The project was supported by the National Natural Science Foundation of China (32122021, 32370452 and 32225009); the National Key R&D Program of China (2022YFC3400300), New Cornerstone Investigator Program to WW and the 1000 Talent Project of Shaanxi Province to K. W. and Q. Q, and the Fundamental Research Funds for the Central Universities.

## Author contributions

K. W., Q. Q. Y. L., and W. W. designed this project and research aspects. C. F., B.W. and M. H. performed sample collection. T. X., F. Z., D. F. and J. H. contributed to sequencing library construction for CUT&Tag and to dual-luciferase assay. M. H. performed the scRNA-seq and bulk RNA-seq data analysis for white spotted bamboo shark, J. Z. conducted the search for CNE. Y. L. conducted the remaining data analysis. C. Z., W. X., Z. L. P. X. provided valuable suggestions for the study. Y. Z. and Z. Z. contributed to the experimental components of this project. Y. L. and K. W. contributed to figure design. Y. L., K. W., Q. Q. and W. W. wrote the manuscript. K. W. and W. W. amended it.

## Data availability

All sequencing data and genome assemble have been deposited in the National Center for Biotechnology Information (NCBI) database (PRJNA1026724).

## Extend Figures

**Extended Data Figure 1.**
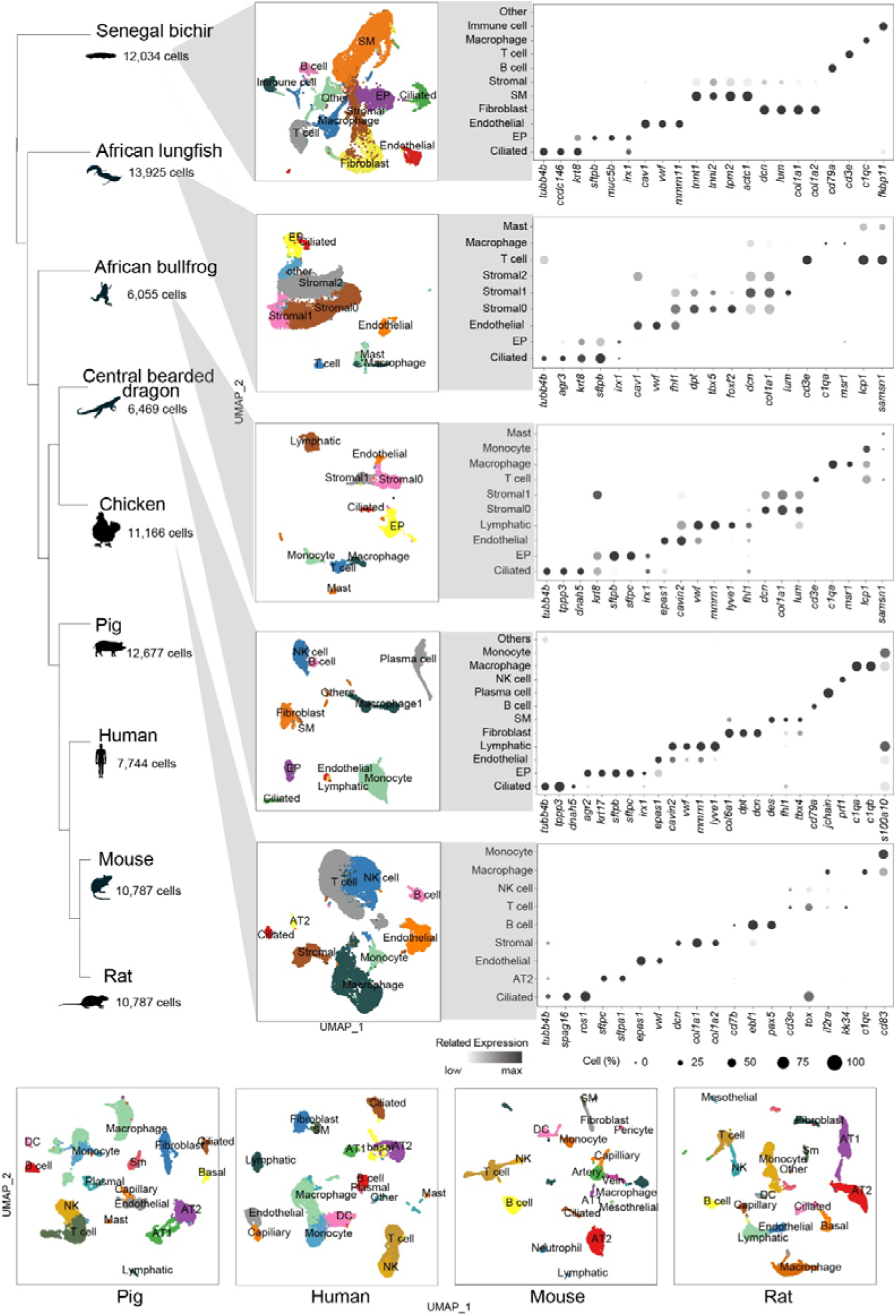
Comparative analysis of cell types and gene expressions across vertebrate species using single-cell RNA sequencing data. This figure displays a phylogenetic tree of various species (from Senegal bichir to rat), with corresponding cell counts and UMAP plots showing cell type clustering. Each species’ plot is color-coded to represent different cell types, including immune, stromal, and epithelial cells. Adjacent to the UMAP plots, a dot plot heatmap illustrates gene expression patterns across cell types, with dot size indicating the percentage of cells expressing each gene and color intensity showing expression levels. The UMAP plots for pig, human, mouse, and rat are provided at the bottom. This visualization enables cross-species analysis of cell compositions and gene expression, highlighting both conserved and divergent features in vertebrate evolution.

**Extended Data Figure 2.**
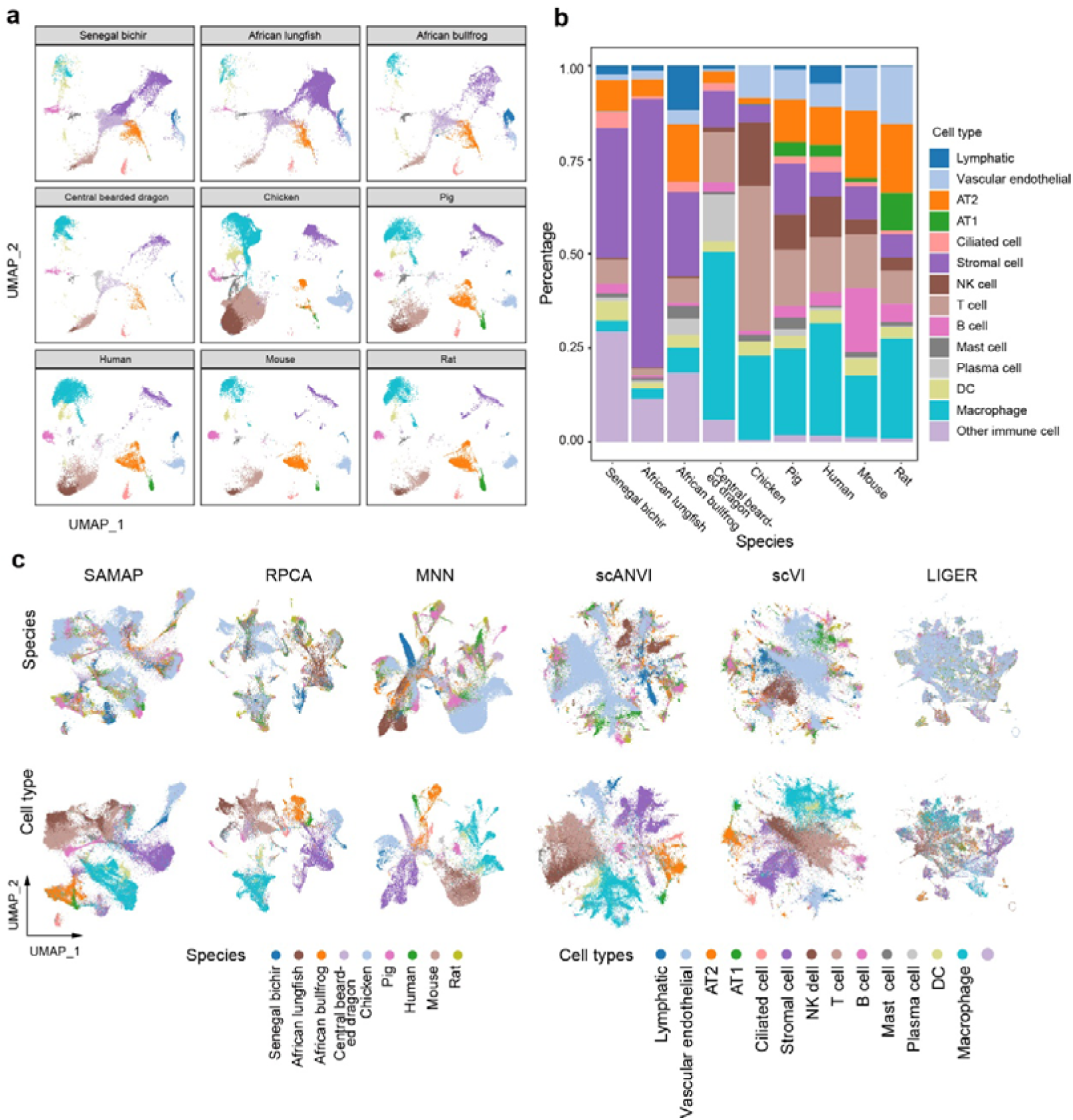
Cross-species comparison and integration of lung cells from nine vertebrate species. **a,** UMAP plots showcasing lung cell distributions for each species, derived from merged Canonical Correlation Analysis (CCA) coordinates. Species range from Senegal bichir to rat, illustrating evolutionary diversity. **b,** Stacked bar chart quantifying the proportional composition of cell types across species. This visualization highlights interspecies variations in lung cellular makeup, from lymphatic and endothelial cells to various immune cell populations. **c,** Comparative UMAP plots demonstrating the integration efficacy of six different computational methods (SAMAP, RPCA, MNN, scANVI, scVI, and LIGER) on lung cells from all nine species. The upper row is color-coded by species origin, while the lower row is color-coded by cell types as annotated from CCA results.

**Extended Data Figure 3.**
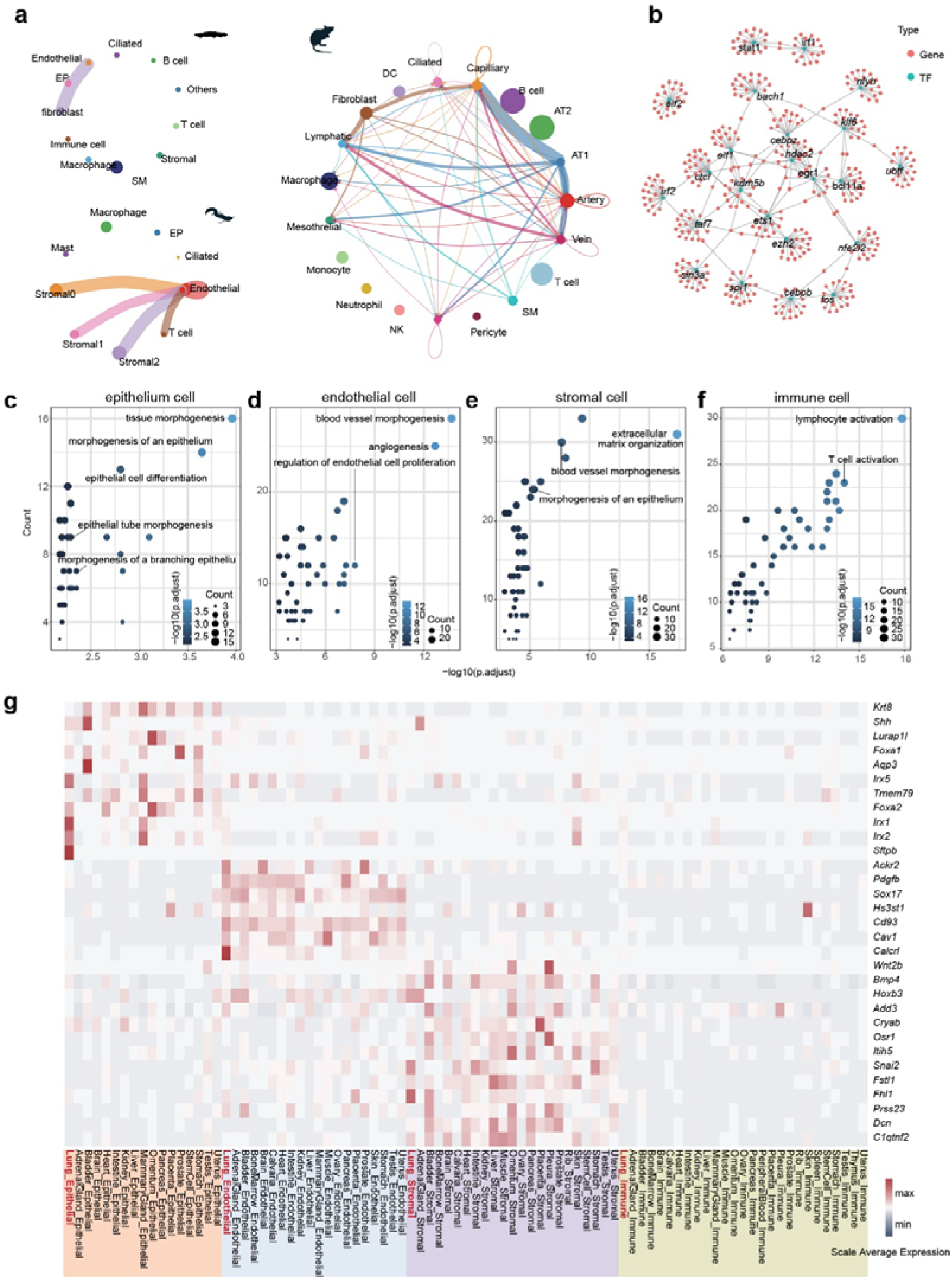
Conservation of cellular communication, transcription factor networks, and gene expression in vertebrate lungs. **a,** Comparative VEGF signaling networks in Senegal bichir, African lungfish, and mouse lungs, derived from CellChat analysis. Circle sizes indicate cell type proportions, while edge widths represent intercellular communication probabilities. This visualization highlights conserved signaling patterns across evolutionarily distant species. **b,** Co-expression network of 23 conserved lung transcription factors (blue nodes) and their partial target genes (red nodes). This network illustrates the complex regulatory relationships governing lung cell identity and function across species. **c-f,** Gene Ontology (GO) term enrichment analysis for conserved genes expressed in lung cell populations across nine vertebrate species. Bubble plots show enriched biological processes for (c) epithelium, (d) endothelial, (e) stromal, and (f) immune cells. Bubble size corresponds to gene count, while color intensity represents statistical significance (-log10 adjusted p-value). **g,** Heatmap depicting the expression patterns of lung-specific genes across various mouse tissue cell populations. Rows represent individual genes, columns represent cell types, and color intensity indicates scaled average expression levels.

**Extended Data Fig. 4.**
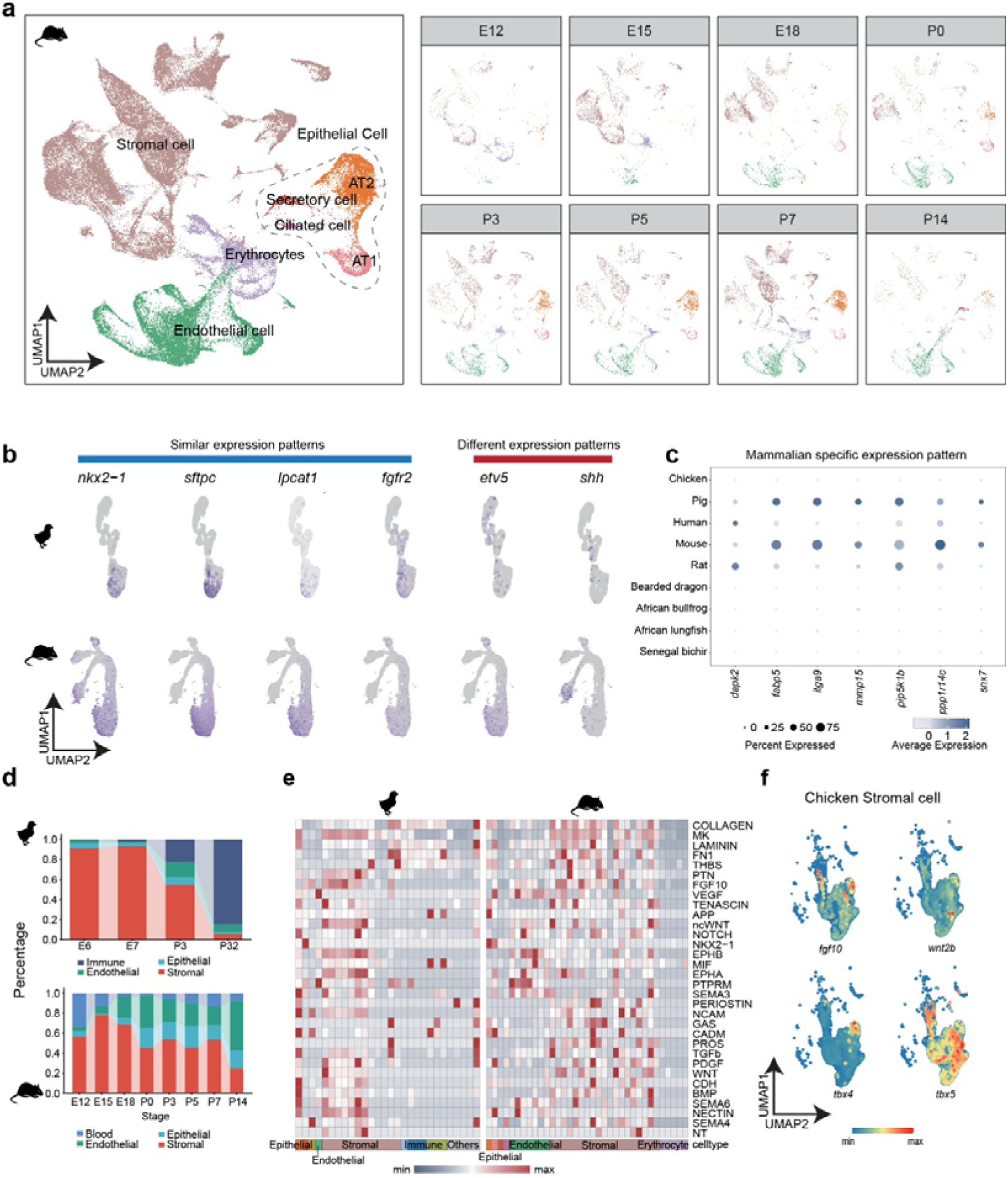
The shared and diverged gene expression pattern between mouse and chicken. **a,** UMAP clustering diagram of mouse lung development. The left plot shows the overall cell type distribution, with major populations such as stromal cells, epithelial cells, endothelial cells, and specialized cell types (AT1, AT2, secretory, and ciliated cells) clearly demarcated. The right series of plots demonstrate the temporal progression of lung cell populations from embryonic day 12 (E12) through postnatal day 14 (P14), showcasing the dynamic changes in cellular composition during development. **b,** Similar and diverged gene expression patterns in epithelial cells between mouse (top row) and chicken (bottom row) during lung development. The left four genes (*nkx2-1*, *sftpc*, *lpcatl1*, *fgfr2*) exhibit similar expression patterns across both species, indicating conserved developmental processes. In contrast, the right two genes (*etv5*, *shh*) show divergent expression patterns, suggesting species-specific adaptations in lung development. **c,** Part of genes with mammalian-specific expression patterns during lung development. The dot plot compares expression levels across various species, from chicken to rat, emphasizing genes that show higher expression or prevalence in mammalian lungs. **d,** The changing proportions of major cell types during lung development in both chicken (top) and mouse (bottom). Notably, the proportion of stromal cells decreases over time in both species, indicating a conserved developmental trend. **e,** Heatmap of 32 shared signaling pathways in stromal cells between mouse and chicken. **f,** The expression of key genes in chicken stromal cells that are known to be highly expressed in mouse lung stromal cells during development.

**Extended Data Figure 5.**
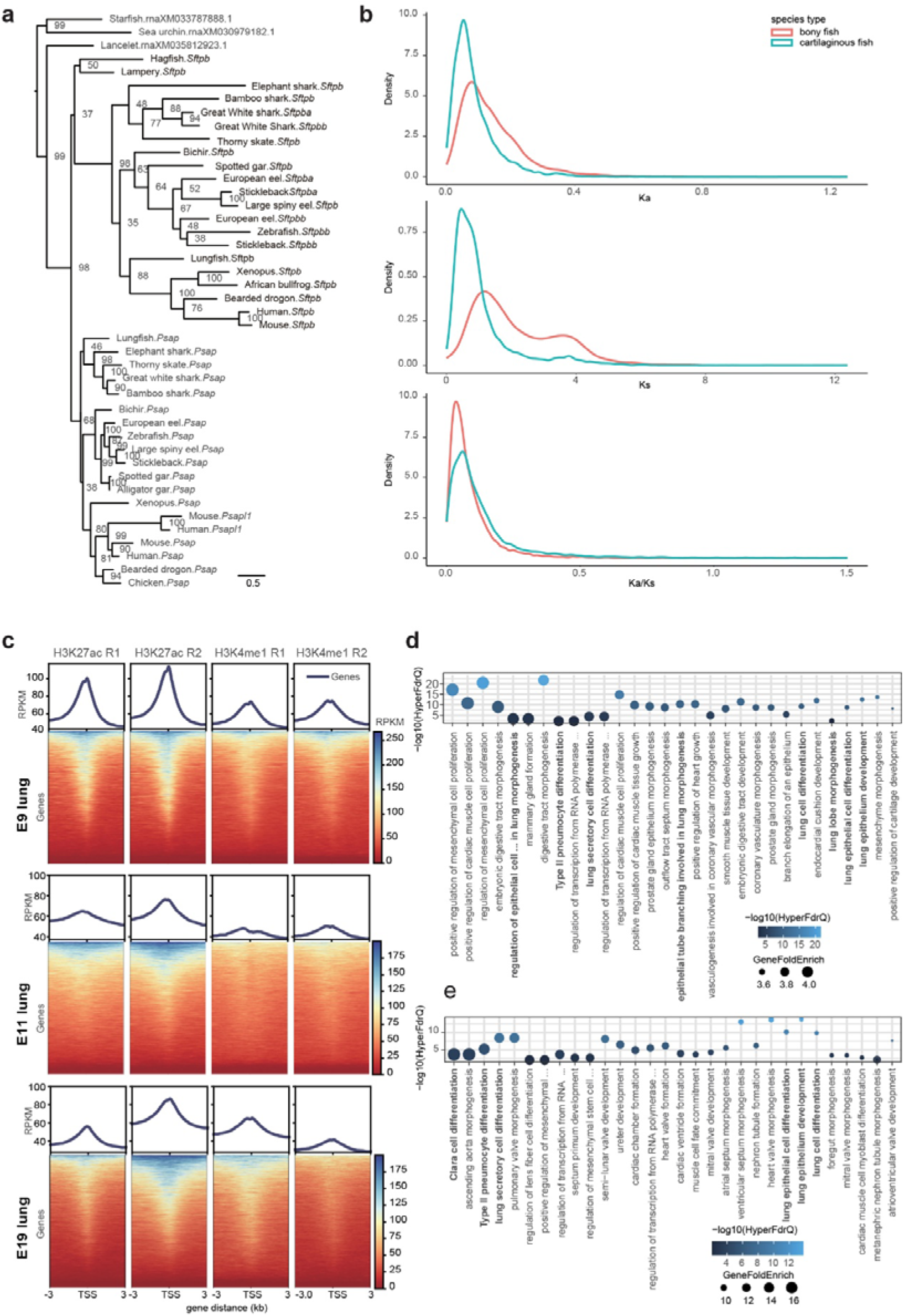
Evolution of lung-related genes in cartilaginous fish and analysis of lung-associated enhancers. **a,** Bayesian phylogenetic tree depicting the evolutionary history of the *sftpb* gene across various species. Numbers at each node represent posterior probabilities (as percentages), indicating the level of support for each branching event. **b,** Comparison of evolutionary rates for lung-related genes between cartilaginous and bony fish. The plots show the distribution of Ka (nonsynonymous substitution rate), Ks (synonymous substitution rate), and Ka/Ks ratio, all calculated relative to the ancestral sequence. **c,** Heatmaps showing the distribution of CUT&Tag signals for histone modifications H3K27ac and H3K4me1 around gene transcription start sites (TSS) in chicken embryonic lungs at 9, 11, and 19 days of development. The panels display signal intensity from 3 kb upstream to 3 kb downstream of the TSS. Blue color indicates high coverage depth, suggesting active regulatory regions. **d,** GREAT (Genomic Regions Enrichment of Annotations Tool) analysis results for all conserved noncoding elements (CNEs) with putative lung regulatory activity. **e,** GREAT enrichment results specifically for CNEs with lung regulatory activity that originated in bony fish.

**Extended Data Figure 6.**
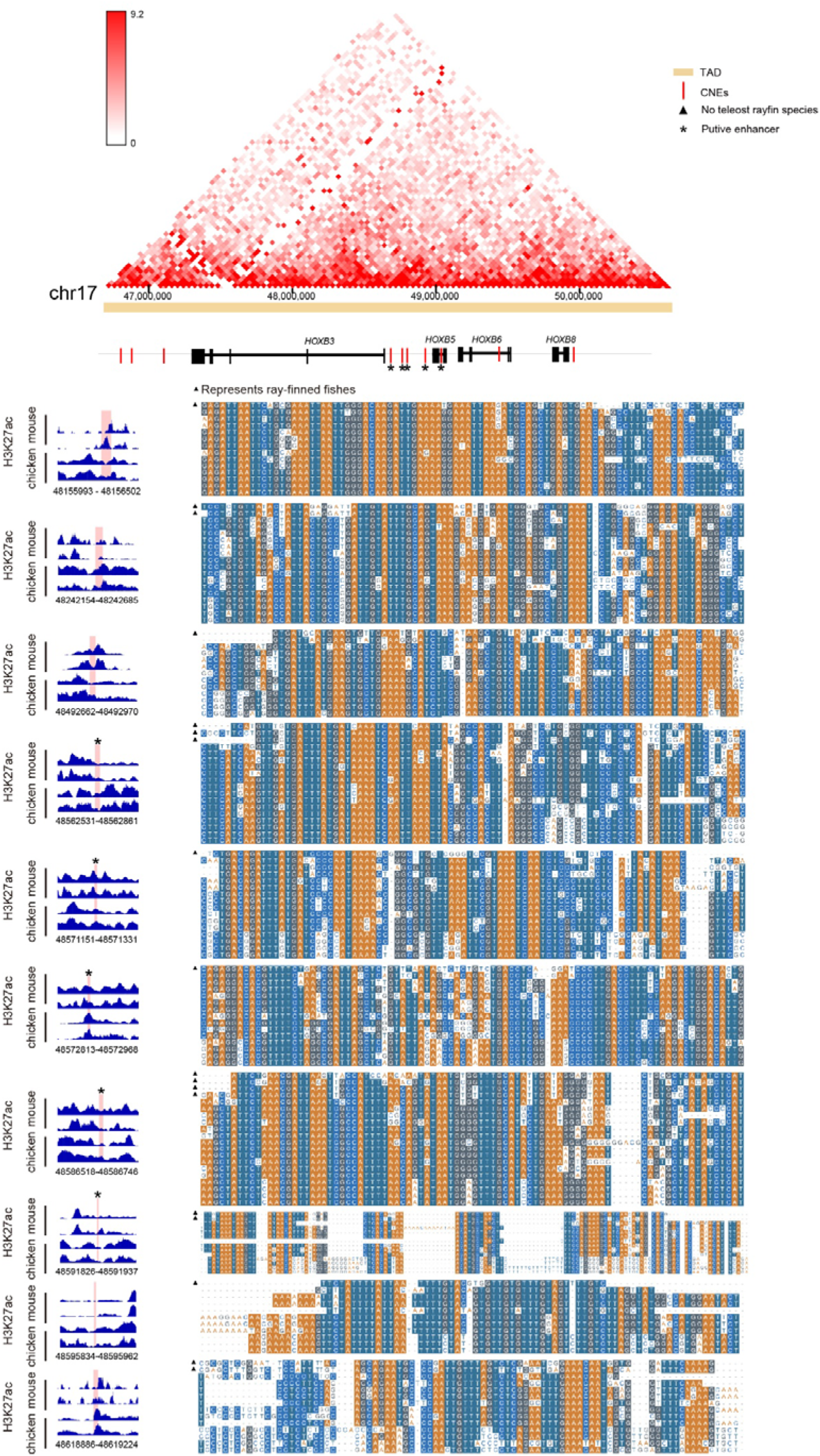
Analysis of 10 lung-regulatory Conserved Noncoding Elements (CNEs) unique to bony fishes within the HOXB gene cluster. Top panel: Hi-C interaction map of the HOXB locus in human lung tissue. The triangular heatmap displays chromatin interaction frequencies, with darker red indicating stronger interactions. This visualization helps identify topologically associating domains (TADs) and potential long-range regulatory interactions. Middle panel: Gene structure and CNE location map of the HOXB cluster (using hg38 as reference). The horizontal bars represent genes, while vertical red lines indicate the positions of CNEs. Black stars beneath some CNEs denote putative enhancers. Bottom panel: Detailed view of 10 CNEs showing: Left: H3K27ac activity profiles in mouse and chicken across various developmental stages and tissues, represented by blue peaks. H3K27ac is a marker of active enhancers, providing evidence for the regulatory potential of these CNEs. Right: Sequence conservation analysis across multiple species. Each row represents a species, with blue boxes indicating sequence conservation and white spaces showing gaps or divergence. Species marked with triangles represent ray-finned fishes, while unmarked species are lobe-finned fishes.

**Extended Data Figure 7.**
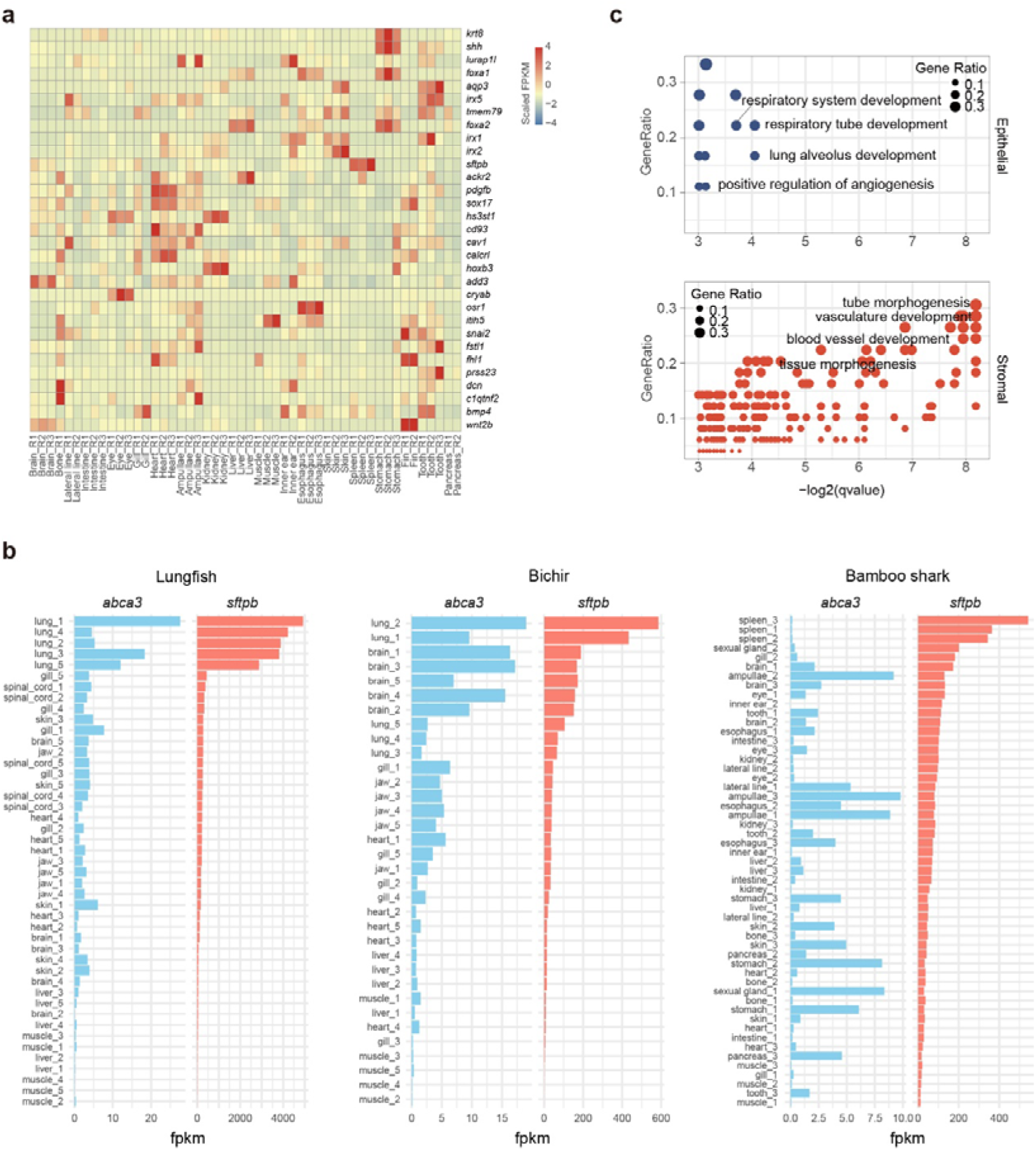
Expression patterns of lung-related genes in cartilaginous fish and their evolutionary changes across vertebrates. **a,** Heatmap showing the expression levels of selected lung related genes (lung-specific genes) in various tissues of the bamboo shark. **b,** Bar plots comparing the expression levels (in FPKM) of two key lung-specific genes, *Sftpb* and *Abca3*, across different tissues in three species: African lungfish, Senegal bichir, and bamboo shark. The plots reveal a degree of co-expression of these genes in lungfish and bichir, particularly in lung tissues, while showing no apparent correlation in the bamboo shark. **c,** Gene Ontology (GO) enrichment analysis of genes that became highly expressed in lung epithelial cells during the terrestrial transition of vertebrates. The upper panel shows enriched terms related to epithelial development, while the lower panel highlights terms associated with stromal development. The size of each dot represents the gene ratio, while the color intensity indicates statistical significance (-log2(q-value)).

**Extended Data Figure 8.**
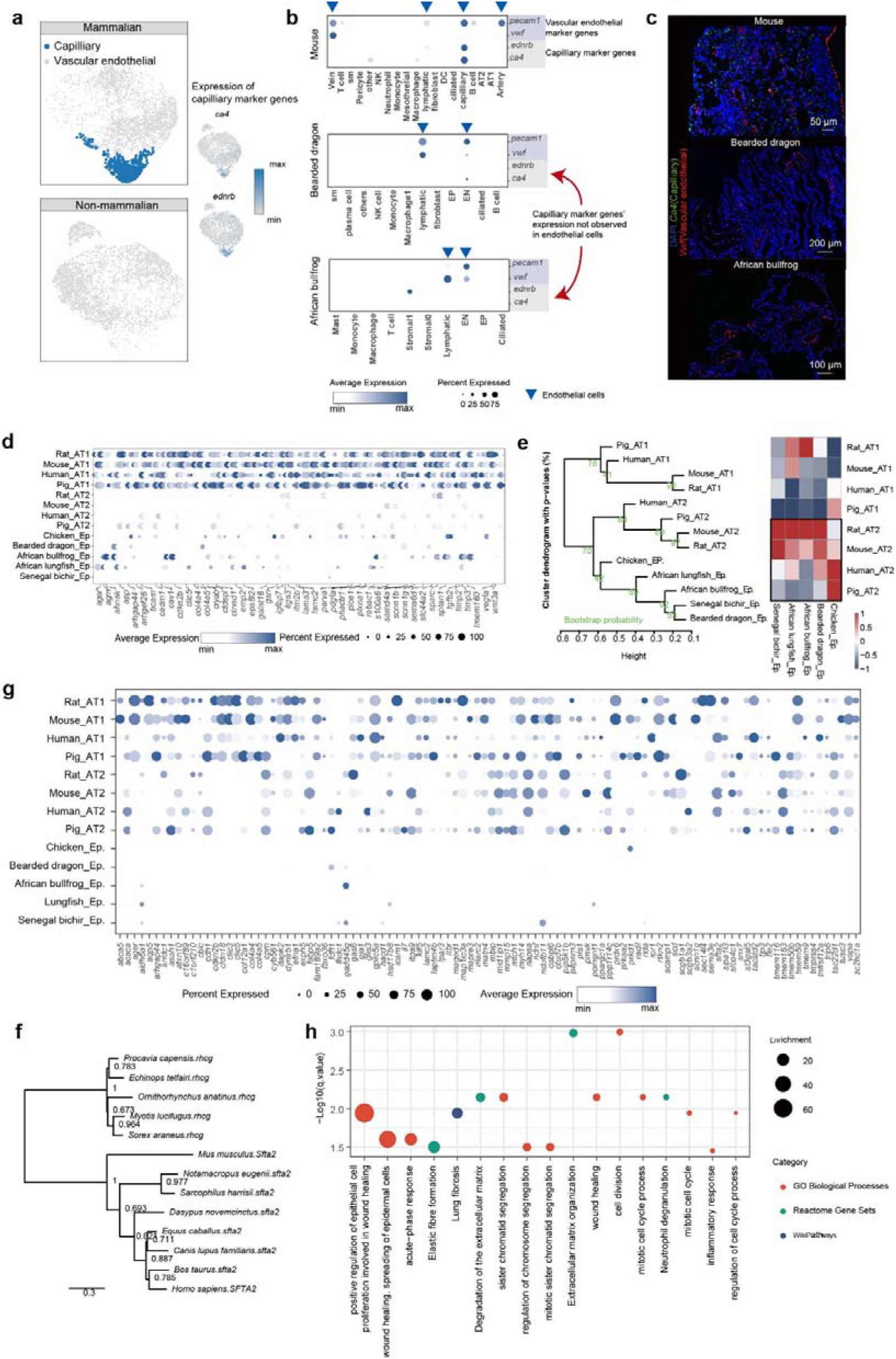
Mammalian-specific adaptations in lung structure and function. **a,** UMAP plots illustrating the further specialization of mammalian lung endothelial cells, particularly highlighting a distinct population of lung capillary cells not identified in the five non-mammalian species studied. The upper panel shows mammalian lung endothelial cells, while the lower panel represents non-mammalian species. **b,** Dot plots demonstrating that capillary marker genes are not prominently expressed in lung endothelial cells of non-mammalian species. Blue triangles indicate endothelial cells in each species. The plot compares gene expression across different cell types and species. **c,** RNA in situ hybridization images of *Ca4* (capillary endothelial cell marker) and *Vwf* (vascular endothelial cell marker) in lung sections from mouse, bearded dragon, and African bullfrog. The results indicate that *ca4* expression is not detectable in the lungs of these two non-mammalian species, supporting the findings from (b). **d,** Heatmap showing genes that are highly expressed in AT1 cells relative to AT2 cells. These genes are also found to be upregulated in mammalian AT1 cells compared to non-mammalian respiratory epithelial cells. **e,** Clustering relationships among AT1, AT2, and respiratory epithelial cells. The left panel shows a phylogenetic tree constructed using expression data, while the right panel represents a similarity matrix. **f,** Phylogenetic tree illustrating the origin of the *sfta2* gene from a duplication of the *rhcg* gene. **g,** Dot plot of genes highly expressed in mammalian epithelial cells compared to non-mammalian respiratory epithelial cells. This includes numerous genes related to lung surfactant, such as *napsa*, *aqp5*, and *prdx6*. Of particular note are the klf5 transcription factor and genes associated with lung surfactant regulation, including *ppargc1a*, *cdh1*, *glis3*, and *prkaa2*. **h,** Enrichment analysis of genes upregulated in the lungs of *sfta2* knockout homozygous mice. The plot shows enriched GO terms, ReactomeDB pathways, and WikiPathways, with dot size indicating gene count and color representing significance.

